# p53 and TIGAR promote redox control to protect against metabolic dysfunction-associated steatohepatitis

**DOI:** 10.1101/2024.05.20.594983

**Authors:** Celine I Wittke, Eric C Cheung, Dimitris Athineos, Nicola Clements, Liam Butler, Mark Hughes, Vivienne Morrison, Dale M Watt, Karen Blyth, Karen H Vousden, Timothy J Humpton

## Abstract

*TP53* is a potent tumour suppressor that coordinates diverse stress response programmes within the cell. The activity of p53 is frequently context and cell type-dependent, and ranges from pro-survival activities, including the implementation of transient cell cycle arrest and metabolic rewiring, through to cell death. In addition to tumour suppressor functions, p53 also has established roles in the pathological response to stress that occurs during tissue damage and repair, including within the liver. Metabolic dysfunction-associated steatohepatitis (MASH) is a major driver of hepatocellular carcinoma (HCC), but our understanding of the molecular determinants of MASH development remains incomplete.

Here, using a p53 reporter mouse, we report early and sustained activation of hepatic p53 in response to an obesogenic high fat and high sugar diet. In this context, liver-specific loss of p53 accelerates the progression of benign fatty liver disease to MASH that is characterised by high levels of reactive oxygen species (ROS), extensive fibrosis, and chronic inflammation. Using an *in vitro* culture system, we show that p53 functions to control ROS and protect against the development of MASH, in part through induction of the antioxidant gene TP53-induced glycolysis and apoptosis regulator (TIGAR). Our work demonstrates an important role for the p53-TIGAR axis in protecting against MASH, and identifies redox control as an essential barrier against liver disease progression.

## INTRODUCTION

Liver cancer (Hepatocellular carcinoma (HCC)) is a common and lethal disease with limited treatment options and increasing prevalence^1, 2^. HCC arising from metabolic dysfunction-associated steatohepatitis (MASH, formerly called non-alcoholic steatohepatitis (NASH)^3^) represents the fastest growing fraction of HCC mortality globally and has become the leading cause of HCC in some western countries, replacing viral hepatitis^2, 4, 5^. Liver disease progression from benign metabolic dysfunction-associated steatotic liver disease (MASLD, formerly called non-alcoholic fatty liver disease (NAFLD)^3^) to diet-induced MASH is strongly linked with the consumption of a high fat and high sugar ‘Western’ diet as well as with features of metabolic syndrome (MetS), including obesity, insulin resistance, hypertension, and hyperlipidaemia^6–9^. Of concern, MASH has especially high prevalence in MetS patients with co-morbid type 2 diabetes mellitus (T2DM). Disease progression is particularly rapid in this cohort, and the development of HCC is more common^10^.

Characteristics of MASH include chronic inflammation, fibrosis, and high levels of reactive oxygen species (ROS), which overlap with hallmarks of cancer^8, 9, 11–13^. This connection is reinforced by data from MASH patients, where cancer-promoting and tumour-suppressive pathways are active, including the classic tumour suppressor protein TP53 (p53)^14–21^. Consistent with these observations, p53 has also been implicated in the aetiology of MetS more broadly. The p53 proline 72 to arginine (P72R) polymorphism, for example, has been reported to promote metabolic dysfunction in a humanised mouse model^22^ and is associated with increased T2DM sequelae in humans^23^.

The p53 transcription factor coordinates a diverse cellular stress response to balance adaptation and cell survival against senescence and cell death^24–28^. As in many tissues, both unrestrained activation of p53 and the absence of p53 in the liver can be lethal^29, 30^. Outside of these severe cases, however, context is an important mediator of p53 activity in the liver. On the one hand, undue p53 pathway activation has been shown to impede liver regeneration, promote fibrosis, and support HCC development following chronic liver damage^31–35^. p53-mediated apoptosis has also been reported to promote MASH in a nutritional stress-induced methionine and choline-deficient (MCD) diet model^36^. Even so, p53 can also exert clear hepatoprotective activity. During damage-induced liver regeneration, for example, p53 has also been shown to limit inflammation and fibrosis, protect against fatty-acid induced apoptosis, act to maintain mitotic fidelity, and promote detoxification of damaging lipid peroxidation ROS^37–43^.

Redox control is an important aspect of p53 function more broadly and is commonly coordinated by downstream antioxidant effector proteins including TP53-inducible glycolysis and apoptosis regulator (TIGAR), amongst many others^25^. TIGAR is a fructose-2,6-bisphosphatase that acts in this capacity to limit accumulation of fructose-2,6-bisphosphate, a potent allosteric activator of phosphofructokinase-1 (PFK-1) and glycolytic activity more generally^44, 45^. In doing so, TIGAR activation dampens PFK-1 activity, thereby reducing glycolytic flux, a response that has been shown to promote the generation of nicotinamide adenine dinucleotide phosphate (NADPH) via increased utilisation of the oxidative pentose phosphate pathway^44–46^. Although it is generally cytoplasmic, TIGAR can also localise to the mitochondria where it exerts additional antioxidant activity that is independent of its fructose-2,6-bisphophatase activity^47, 48^. Functionally, TIGAR has been shown to support intestinal regeneration and promote tumourigenesis, in part by enhancing the detoxification of peroxidised lipids and keeping ROS levels low^49^.

Here, we investigate the importance of p53 activity as a mediator of diet-induced liver disease progression. Our work demonstrates an important role for p53-mediated redox control in protecting against the development of MASH *in vivo*. We have further identified TIGAR as an important mediator of protective p53 activity in this context and provide evidence that antioxidant therapy can ameliorate features of MASH.

## RESULTS

### Liver p53 is engaged in response to a high fat and high sugar ‘Western’ diet

In humans, increased expression of either *TP53* or the p53 target gene *CDKN1A/p21* (p21) is correlated with MASH, liver fibrosis, and T2DM^20, 21^. These observations suggest a potential role for p53 in mediating the aetiology of diet-induced liver disease. We have previously utilised a p53 reporter mouse to non-invasively monitor the p53 pathway after total body gamma-irradiation and during hepatotoxin-mediated liver regeneration^50^. In p53 reporter mice, active p53 induces expression of the near-infrared fluorescent protein iRFP713 (iRFP) that is linked to the synthetic PG13 p53-responsive promoter (Fig. S1A). The resulting fluorescence can be monitored non-invasively and longitudinally^50^.

To explore the dynamics of p53 activity during diet-induced liver disease, male and female p53 reporter mice were shifted onto either a purified high fat and high sugar (HFHS) ‘Western’ diet containing 59% energy from fat, 15% energy from protein, 25% energy from carbohydrates, and rich in sucrose^51^ or onto a purified control diet containing 10% energy from fat, 14% energy from protein, 76% energy from carbohydrates, and reduced sucrose content. Mice were maintained on either diet for a period of up to 300 days and regularly monitored for weight gain and iRFP expression (Figs. 1A-F and S1B). As expected, both male and female mice exhibited weight gain on the HFHS diet relative to mice on the control diet (chow) within the first 50 days (Fig. 1E/F). Consistent with reports linking p53 activity with MASH^20, 21, 36^, we observed robust signal from the p53 reporter in both male and female mice by the end of our study (Figs. 1A-D and S1B). Interestingly, there was also significant and sustained activation of the p53 reporter within male mice beginning much earlier, starting at 100 days of the HFHS diet and increasing thereafter (Fig. 1A/B). At this timepoint, although HFHS-fed mice of both sexes had gained significant weight, they did not exhibit significant changes in oral glucose tolerance compared with mice on the control diet (Fig. S1C), suggesting that diet-induced MetS was still in the early stages. Within female HFHS-fed mice, even though initial weight gain occurred similarly rapidly, induction of the p53 reporter was a much later event than in male HFHS-fed mice—at 200 days of the HFHS diet—and occurred with significantly lower fold iRFP reporter induction compared with male mice (Figs. 1C/D and S1D).

**Figure 1:**
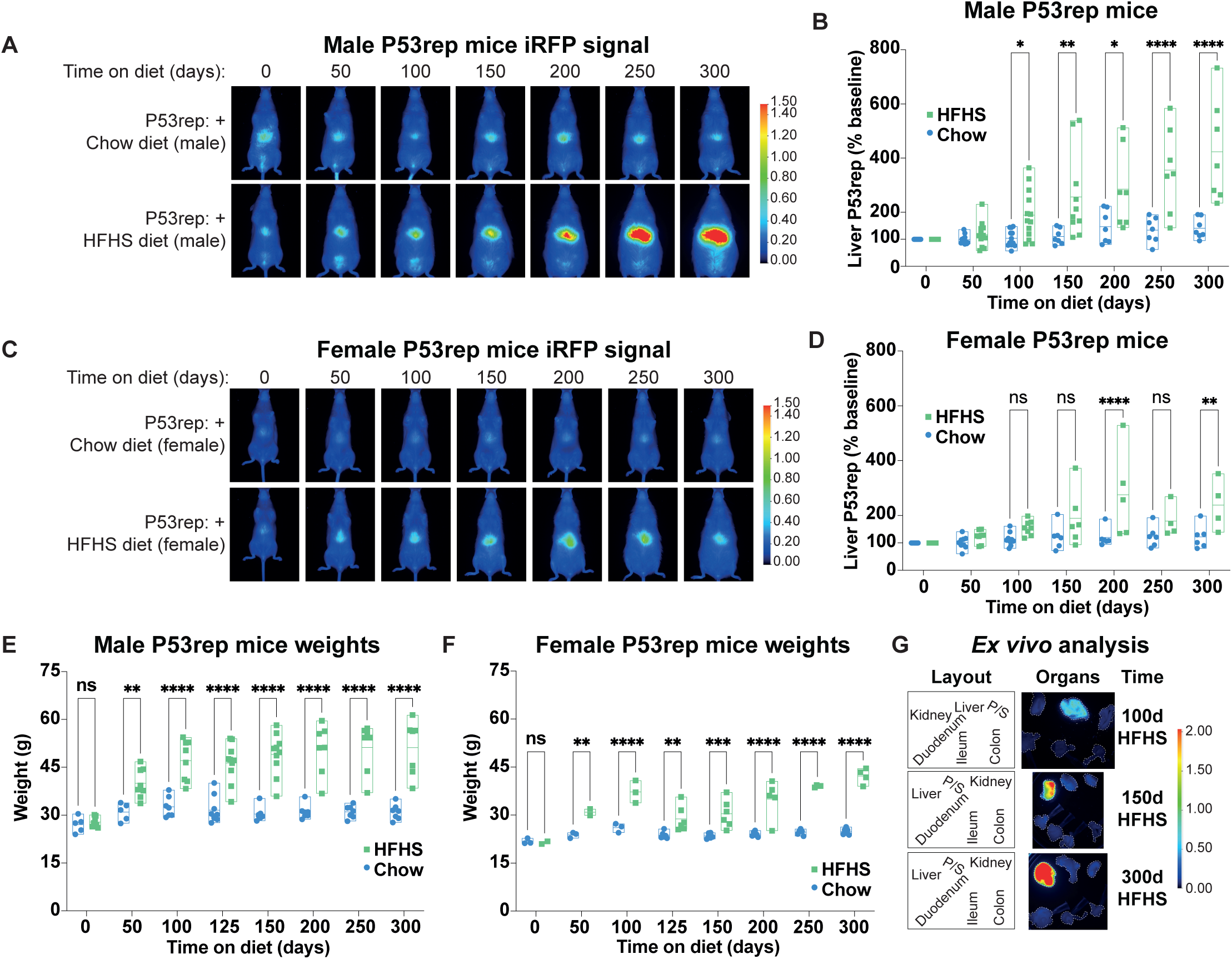
Liver p53 is engaged in response to an obesogenic high fat high sugar diet. A/B. Male P53rep mice were given obesogenic high fat high sugar (HFHS) diet at 65-75 days of age or put onto an imaging-suitable control diet (chow) and imaged after the indicated days. Representative images (A) and P53rep signal quantification (B) normalised to the initial liver signal identified per mouse. N=12 chow and N=13 HFHS mice at 0d, 50d, 100d; N=7 chow and N=10 HFHS at 150d; N=7 chow and N=7 HFHS at 200d, 250d, 300d. Data presented as mean +/- range with individual data points (B). Data analysed using a Two-way ANOVA with Holm-Sidak’s multiple comparisons test and multiplicity-adjusted p- values. *p<0.05, **p<0.01, ****p<0.0001. C/D. Representative images (C) and quantifications (D) of female mice on same diet and imaging regime as in (A/B). N=9 chow and N=8 HFHS mice for 0d, 50d, 100d; N=6 chow and N=6 HFHS for 150d, N=6 chow and N=5 HFHS at 200d; and N=6 chow and N=4 HFHS for 250d and 300d. Data presented and analysed as (B). ns- not significant, **p<0.01, ****p<0.0001. E. Weight (g) of male HFHS-fed and control diet mice from (A/B). Timepoints prior to 150d reflect a subset of the mice. N=5 chow and N=8 HFHS for 0d and 50d, N=7 chow and N=8 HFHS mice for 100d, N=9 chow and N=10 HFHS for 125d, N=7 chow and N=10 HFHS mice for 150d, and N=7 mice per condition for 200d, 250d, and 300d timepoints. Data presented and analysed as (A/B). ns- not significant, **p<0.01, ****p<0.0001. F. Weight (g) of female HFHS-fed and control mice as in (E). Timepoints prior to 125d reflect a subset of mice. N=3 chow and N=2 HFHS mice for 0d and 50d, N=3 mice each for 100d, N=6 chow and N=6 HFHS for 125d and 150d, N=6 chow and N=5 HFHS for 200d, N=6 chow and N=4 HFHS at 250d and 300d. Data presented and analysed as (A/B). ns- not significant, **p<0.01, ***p<0.001, ****p<0.0001. G. Representative images of *ex vivo* P53rep iRFP signal in tissues after 100d, 150d, or 300d on diet. Organ layout and LUT intensity profile as shown. Representative of N=3 male mice/timepoint. P/S- Pancreas/Spleen.

Although it is possible to differentiate the spatial distribution of iRFP expression on whole-body scans, for example between liver- and gut region-specific signals, organ overlap within a region of interest can sometimes obfuscate the source of p53 reporter signal within the two-dimensional images. *Ex vivo* analysis of organs is unambiguous, however, and confirmed that p53 induction was restricted to the liver in HFHS-fed mice sampled throughout the duration of the study (after 100d, 150d, and 300d of HFHS diet) (Fig. 1G).

Together, these findings reveal that p53 induction is an early and sustained feature of the hepatic response to chronic consumption of a high fat and high sugar ‘Western’ diet. Although male and female mice both induce p53, its induction is particularly rapid and robust in male mice *in vivo*.

### Liver-specific loss of p53 accelerates ‘Western’ diet-induced MASH

Within the liver, p53 activity can be both protective and damaging, seemingly determined in part by the extent and duration of underlying liver damage^34^. To explore potential functional roles for p53 activity during diet-induced liver disease, we created cohorts of mice harbouring liver-specific deletion of *Trp53* (*p53*) (*Albumin-Cre+*; *Trp53^FL/FL^* mice) and proceeded to characterise the long-term response to the HFHS diet within this model. Importantly, *Albumin-*Cre*+*; *Trp53^FL/FL^*mice retain wildtype p53 expression in all tissues except the liver. They are therefore not susceptible to developing cancer before approximately 2 years of age, as previously reported^43^, and allow for long-term studies into the effects of p53 loss on liver biology. Based on our findings in Figure 1, we focused on examining the response to the HFHS diet within male mice in these experiments to provide the best signal-to-noise resolution of potential aspects of p53 function.

In *Albumin-*Cre*+*; *Trp53^FL/FL^* male mice, we observed clear signs of MASH after a period of one year on the HFHS diet that remained largely absent in *Trp53* wildtype mice fed the HFHS diet (Figs. 2A-E and S2A-C). Compared with either *Albumin-*Cre*+*; *Trp53^FL/FL^*mice on the control diet or with *Albumin-*Cre*+*; *Trp53* wildtype mice on the HFHS diet, *Albumin-* Cre*+*; *p53^FL/FL^* HFHS diet fed mice exhibited histological features of MASH, including abundant steatosis, hepatocyte ballooning, the accumulation of intracytoplasmic proteins known as Mallory-Denk bodies, hepatic hypertrophy, and inflammatory infiltrates in the liver (Figs. 2A and S2A). MASH within *Albumin-*Cre*+*; *Trp53^FL/FL^* HFHS diet fed mice was further characterised by significantly increased levels of immunohistochemical (IHC) staining for malondialdehyde (MDA), a marker of lipid peroxidation, substantial staining for picrosirius red (PSR), a marker of fibrosis, and a clear increase in hepatic lipogranulomas, consistent with increased chronic inflammation (Figs. 2A-E and S2A). This was not the case in *Albumin-*Cre*+*; *Trp53* wildtype HFHS diet fed mice, where features of MASLD, such as lipid accumulation (Figs. 2A/B and S2A) were also evident, but with significantly lower staining for further markers indicative of MASH, consistent with greater ongoing damage in *Albumin-Cre+*; *Trp53^FL/FL^*HFHS diet fed mice compared with *Albumin-*Cre*+*; *Trp53* wildtype HFHS fed mice.

**Figure 2:**
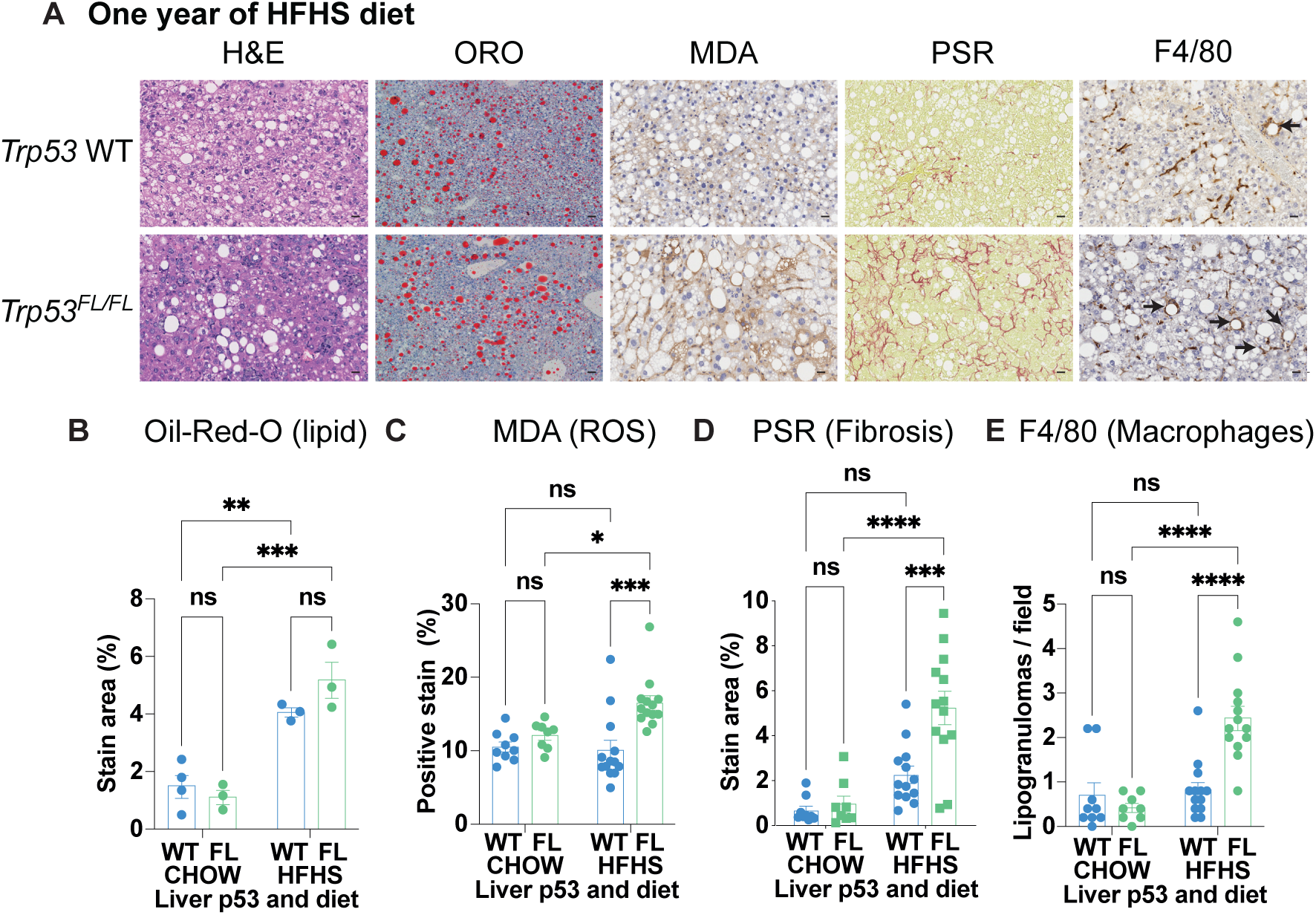
Liver p53 limits damaging ROS, fibrosis, and inflammation associated with MASH. A. Representative liver H&E images, staining for lipid content (oil-red-o / ORO), IHC staining for lipid peroxidation (malondialdehyde/MDA), staining for fibrosis (picrosirius red/PSR), and IHC staining for macrophages (F4/80) in male *Albumin*-Cre+; *Trp53* WT and *Albumin*-Cre+; *Trp53^FL/FL^*(liver-specific deletion of p53) (*p53^FL/FL^*) mice after one year on obesogenic high fat high sugar (HFHS) diet. ORO images are representative of N=4 *Trp53* WT chow-fed mice, N=3 *Trp53^FL/FL^* chow-fed mice and N=3 HFHS-fed mice per genotype; scale bars 40 μm. For the remainder, representative of N=9 *Albumin*-Cre+; *Trp53* WT chow-fed mice, N=8 *Albumin*-Cre+; *Trp53^FL/FL^* chow-fed mice and N=13 HFHS-fed mice per genotype. Scale bars 20 μm. Arrows denote F4/80-positive macrophage-engulfed hepatocytes (lipogranulomas) indicative of chronic inflammation. B-E. Quantification of oil-red-o (ORO) to assess lipid content (B), malondialdehyde (MDA) to assess lipid peroxidation (C), picrosirius red (PSR) to assess fibrosis (D), and F4/80 to assess lipogranuloma formation (E) in male mice after one year on the HFHS diet from (A) compared with aged-matched mice on control diet (chow). N-numbers as in (A). Data presented as mean +/- SEM and analysed using two-way ANOVA with Holm-Sidak’s multiple comparisons test and multiplicity-adjusted p-values. ns- not significant, *p<0.05, **p<0.01, ***p<0.001, ****p<0.0001.

Functionally, plasma levels of alanine transaminase (ALT) and aspartate aminotransferase (AST) enzyme activity were both elevated in *Albumin-Cre+*; *Trp53^FL/FL^* HFHS diet fed mice, consistent with liver damage and compromised liver function in these mice compared to other cohorts at the conclusion of the study (Fig. S2B/C). Even so, overall survival within this time period was not significantly different between *Albumin-Cre+*; *Trp53* wildtype and liver *Trp53*-deficient HFHS diet mice, suggesting a longer latency was necessary for consistent development of MASH-HCC within this HFHS diet model (Fig. S2D). Based on these findings, we concluded that p53 exerts a protective function to oppose MASLD to MASH progression *in vivo*. Although the presence of wildtype *p53* did not alter lipid accumulation resulting from long-term HFHS diet consumption, downstream detrimental effects, including undue lipid peroxidation, fibrosis, and inflammation were significantly reduced—and liver function was maintained—in HFHS diet mice that retained p53 activity.

### Loss of liver p53 is lethal in the Lep^ob/ob^ leptin-deficient genetic model of obesity

Leptin-deficient *Lep^ob/ob^* (*ob/ob*) mice develop progressive liver disease with features of MASLD from an early age that progresses to include evidence of MASH within 20-30 weeks as a consequence of chronic hyperphagia^52, 53^. To examine whether p53 also exerts protective activity against liver disease within this context, we created cohorts of male and female *ob/ob* mice harbouring liver-specific deletion of *Trp53* (*ob/ob; Albumin-*Cre+; *Trp53^FL/FL^* mice) and proceeded to assess liver disease development within this model compared with *Trp53* wildtype *ob/ob; Albumin-*Cre+ mice.

Consistent with published reports^52^, we observed significant steatosis in our *Trp53* wildtype *ob/ob; Albumin-*Cre+ mice but this did not appreciably progress to MASH within the study period (Fig. 3A-C). Unexpectedly, however, the loss of liver *Trp53* was lethal in both male and female *ob/ob; Albumin-*Cre+; *Trp53^FL/FL^* mice (Fig. 3A/B). In comparison to *Trp53* wildtype *ob/ob* mice, where all cohort mice survived to the study endpoint, a median survival of just 319 days was observed in *ob/ob; Albumin-*Cre+; *Trp53^FL/FL^*mice (Fig. 3A/B). Survival within this cohort appeared to be independent of sex, with both male and female mice exhibiting a median survival based on humane clinical endpoint of between 295 days of age (male) and 319 days (female). Histopathological analysis of end-point samples confirmed a preponderance of liver tumours within *ob/ob; Albumin-*Cre+; *Trp53^FL/FL^*mice (8/10 with evidence of liver tumours at necropsy with 5/10 confirmed to have significant tumour burden by histopathology), consistent with a strong liver-related phenotype.

**Figure 3:**
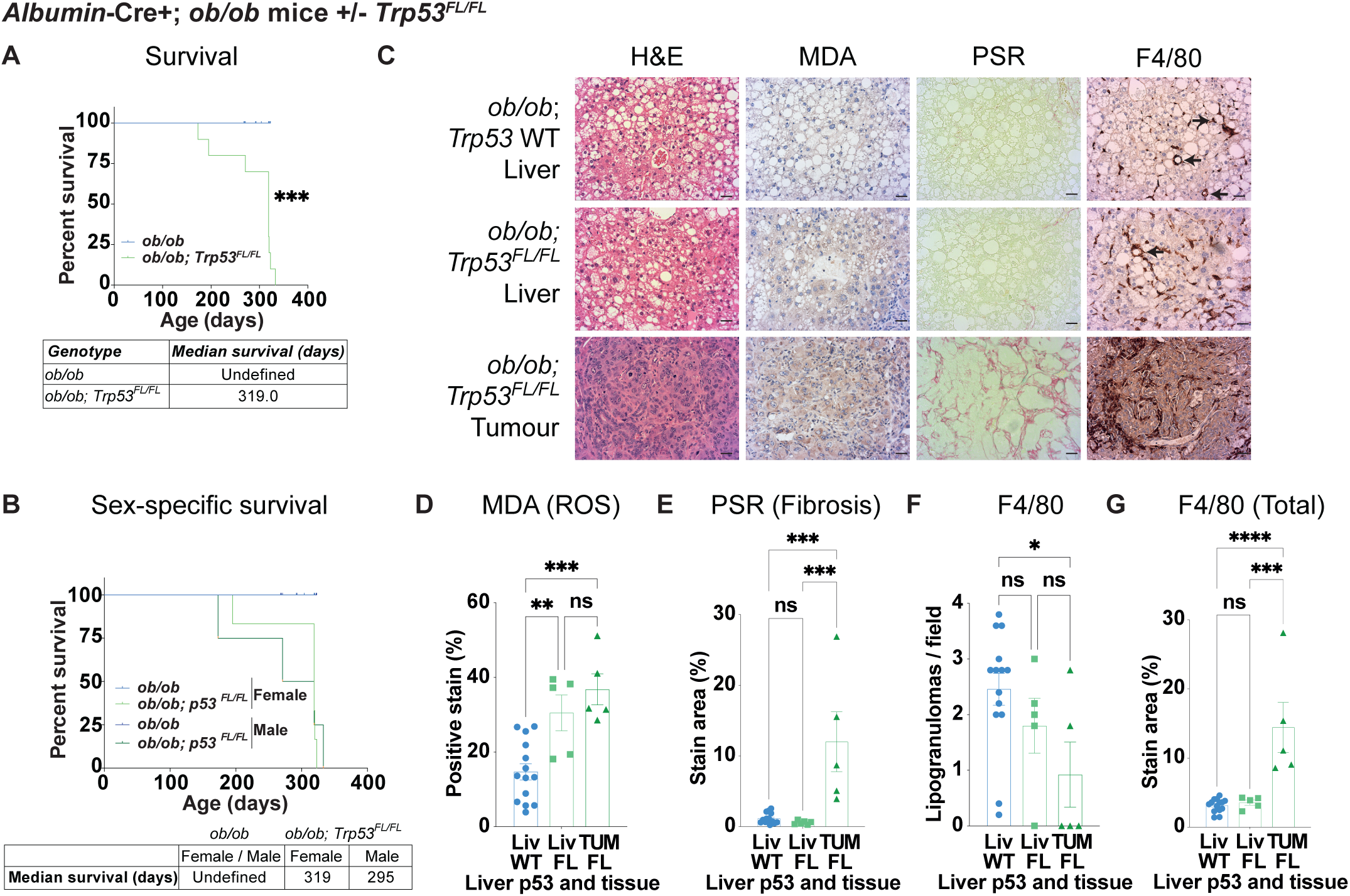
Loss of liver p53 is lethal in leptin-deficient (*Lep^ob/ob^*) obese mice. A. Survival percentages of *Albumin*-Cre+; *Lep^ob/ob^*; *Trp53* WT (*ob/ob*) and *Albumin*-Cre+; *Lep^ob/ob^*; *Trp53^FL/FL^* mice (*ob/ob*; *p53^FL/FL^*). N=8 male and 6 female *ob/ob*; *Trp53* WT mice (14 total) and N=4 male and N=6 female *ob/ob*; *Trp53^FL/FL^* mice (10 total). Data analysed using the Log-rank (Mantel-Cox) test. ***p<0.001. *Ob/ob*; *Trp53^FL/FL^* mice were sampled when reaching endpoint due to clinical signs of moderate severity. Of these, N=5/10 had liver tumours that were confirmed by histology and N=5/10 had small tumours or lacked macroscopic tumour burden, as reflected in downstream analyses. *ob/ob*; *Trp53* WT mice were all healthy at endpoint and sampled at timepoints consistent with the observed survival of *ob/ob*; *Trp53^FL/FL^* mice, in a time range from 269d-323d of age. B. Survival percentages of mice from (A) separated by sex. C. Representative liver H&E images, IHC staining for malondialdehyde (MDA), staining for fibrosis (picrosirius red/PSR), and IHC staining for macrophages (F4/80) in mice from (A). N=14 *ob/ob*; *Trp53* WT mice, N=5 *ob/ob*; *Trp53^FL/FL^*mice without macroscopic tumour on histological analysis and N=5 *ob/ob*; *p53^FL/FL^* mice with tumours (TUM). Macrophage-engulfed steatotic hepatocytes (lipogranulomas) depicted by arrows. Scale bars 20 μm. D-G. Quantification of liver images in C, MDA to assess lipid peroxidation (D), PSR to assess fibrosis (E), F4/80 to analyse lipogranulomas (F), and total F4/80 to assess total macrophage content (G) in *Albumin*-Cre+; *Lep^ob/ob^*; *Trp53* WT (Liv WT) and *Albumin*-Cre+; *Lep^ob/ob^*; *Trp53^FL/FL^* mice (Liv FL and TUM FL) mice at endpoint from (C). N-numbers as in (C). Data presented as mean +/- SEM and analysed using an ordinary one-way ANOVA and Holm-Sidak’s multiple comparisons test and multiplicity-adjusted p-values. ns- not significant, *p<0.05, **p<0.01, ***p<0.001, ****p<0.0001.

As with *Albumin-*Cre+; *Trp53^FL/FL^* mice on the HFHS diet (Fig. 2), liver tissue sections from *ob/ob; Albumin-*Cre+; *Trp53^FL/FL^* mice at endpoint were characterised by elevated IHC staining for MDA (Fig. 3C/D). Liver tumours from *ob/ob; Albumin-*Cre+; *p53^FL/FL^* mice also exhibited extensive fibrosis (Figs. 3C and 3E). Unlike in the HFHS diet model, however, *ob/ob; Albumin-*Cre+; *Trp53^FL/FL^* tumour mice had fewer lipogranulomas than age-matched *Trp53* wildtype *ob/ob* mice, although overall macrophage infiltration was significantly increased (Figs. 3C and 3F/G). These potentially related findings could be due to differences in macrophage response to MASLD/MASH in the *ob/ob* model compared with diet-induced disease, which is in accordance with previous studies^52^. It could also relate to the nature of the tumours observed in *ob/ob; Albumin-*Cre+; *Trp53^FL/FL^* mice, which generally lacked visible steatosis (Fig. 3C). Collectively, these findings suggest that as in diet-induced liver disease, liver p53 acts to limit MASLD to MASH progression in the *ob/ob* leptin-deficient genetic model of obesity *in vivo*.

### P53 engages TIGAR and supports redox control in response to nutrient excess

Comparison of our findings across diet-induced and genetic models of liver disease highlighted diminished redox control as one of the common connecting characteristics. We have previously reported on protective functions of p53-mediated redox control in the liver during regeneration^43^ and sought to further interrogate the possibility of a similar paradigm operating to protect against MASH. To do so, we developed an *in vitro* medium formulation based on Plasmax, a culture medium that was developed to mimic human plasma^54^, that additionally matched the relative proportions of the major fatty acids (steric, palmitic, oleic, linoleic, and linolenic acids) and carbohydrate sources (glucose and fructose) found within our murine HFHS diet. In order to ensure their bioavailability, the fatty acids in our formulation were conjugated to bovine serum albumin (BSA) to improve solubility within medium. For this reason, we utilised Plasmax supplemented with an equivalent amount of fatty acid-free BSA as the control ‘baseline’ formulation for *in vitro* experiments.

To explore the role of p53 in hepatocytes during the response to diet stress more fully *in vitro*, we turned to murine (Hep53.4) and human (HepG2) HCC cell lines that maintain wildtype p53^43, 55^. Consistent with the observations in HFHS diet fed mice, treatment of either cell line with HFHS Plasmax medium for a period of 48 hours increased expression of the potent p53 target genes p21 and MDM2^56^, as well as elevated the levels of TP53-inducible glycolysis and apoptosis regulator (TIGAR)—a key mediator of p53 redox regulation (Fig. 4A-D) ^25, 26^. This response was abrogated in HepG2 cells with *CRISPR*- mediated knock-out of *TP53* (*p53* KO), confirming the role of p53 in the upregulation of TIGAR expression in response to HFHS medium (Fig. 4C/D).

**Figure 4:**
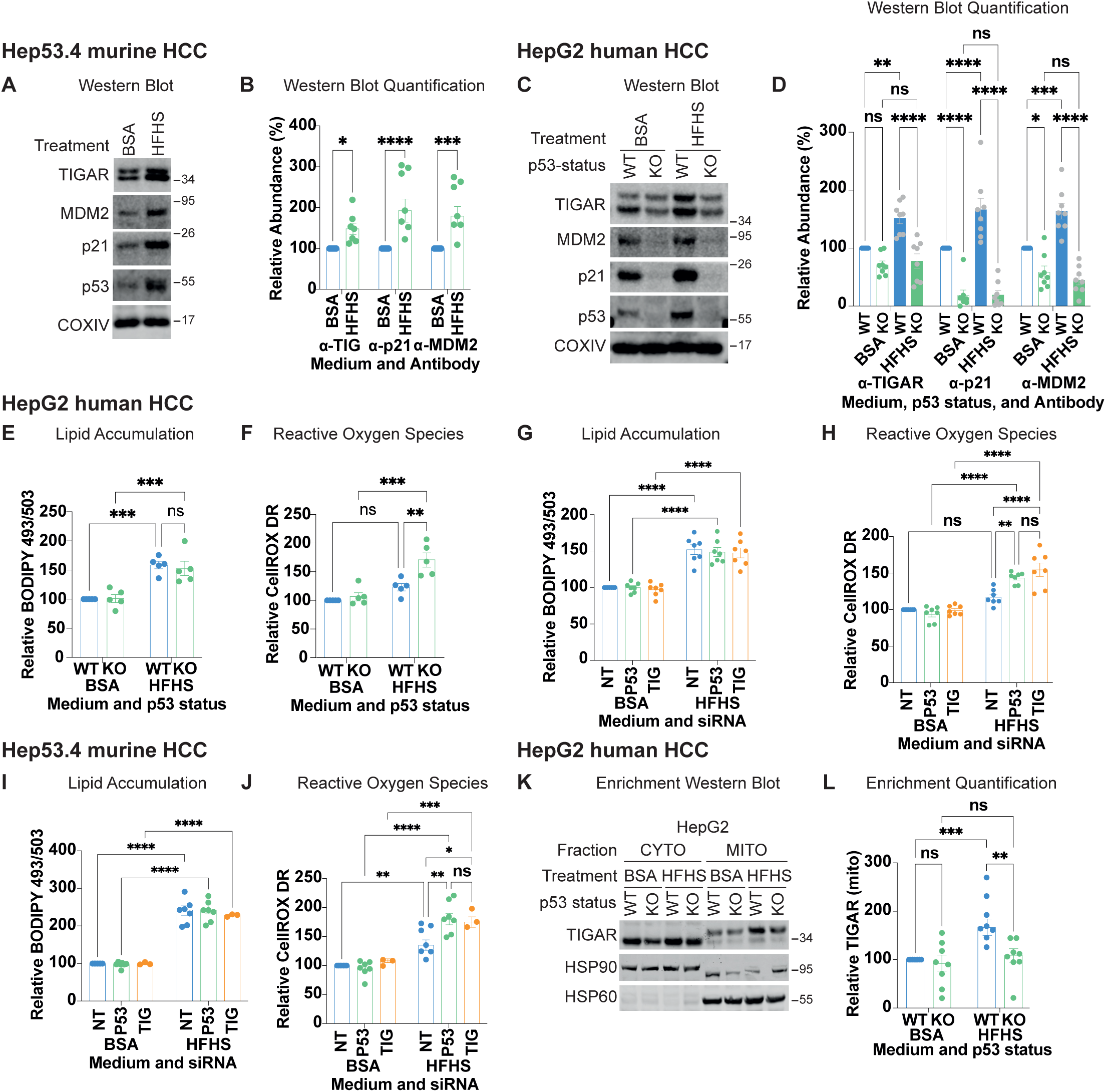
p53 engages TIGAR and supports redox control in response to overfeeding *in vitro*. A/B. Western blot analysis (A) and quantification (B) of TIGAR, MDM2, and CDKN1A/p21 expression in the p53 WT Hep53.4 HCC cell line as cultured in baseline medium supplemented with BSA (BSA) or the high fat high sugar (HFHS) medium formulation for 48 hours. Representative of N=5 independent experiments. COXIV used as loading control. Protein MW ladder (in kilodaltons) as depicted. Data presented as mean +/- SEM with data points and analysed by two-way ANOVA with Holm-Sidak’s multiple comparisons test and multiplicity-adjusted p-values: *p<0.05, ***p<0.001, ****p<0.0001. C/D. Western blot analysis (C) and quantification (D) of TIGAR, MDM2 and CDKN1A/p21 expression in the p53 WT HepG2 HCC and p53-deficient (KO) HepG2 cells generated through CRISPR-mediated p53 targeting as in (A). Representative of N=8 independent experiments. COXIV used as loading control. TIGAR abundance represents sum of bands. Protein MW ladder (in kilodaltons) as depicted. Data presented as mean +/- SEM with data points and analysed by two-way ANOVA with Holm-Sidak’s multiple comparisons test and multiplicity-adjusted p-values: ns- not significant, *p<0.05 **p<0.01, ***p<0.001, ****p<0.0001. E/F. Analysis of lipid content (BODIPY 493/503) (E) and ROS (CellROX Deep Red) (F) via flow cytometry in p53 WT and p53 KO HepG2 cell lines cultured in same media formulations as (A/B). Data from N=5 independent experiments, presented and analysed as (B). ns- not significant, **p<0.01, ***p<0.001. G/H. Analysis of lipid content (BODIPY 493/503) (G) and ROS (CellROX Deep Red) (H) via flow cytometry in p53 WT HepG2 cells with either non-targeting control (NT) or siRNA against *TP53* (P53) or *TIGAR* (TIG) for 96 hours and cultured as (A/B). Data from N=7 independent experiments and presented as median fluorescence intensity (MFI) relative to NT baseline condition per experiment as mean with individual data points per experiment. Data analysed as in (B). ns- not significant, **p<0.01, ****p<0.0001. I/J. Analysis of lipid content (BODIPY 493/503) (I) and ROS (CellROX Deep Red) (J) via flow cytometry in the p53 WT Hep53.4 HCC cell line treated with either non-targeting control (NT) or siRNA directed against *Trp53* (P53) or *Tigar* (TIG) for 72 hours and cultured as (A). Data from N=7 experiments for NT and *TP53* siRNA and N=3 for TIGAR siRNA. Data presented and analysed as in (E/F): ns- not significant, *p<0.05, **p<0.01, ***p<0.001, ****p<0.0001. K. Western blot analysis of TIGAR, MDM2 and CDKN1A/p21 expression in p53 WT and p53-deficient (KO) HepG2 cells as cultured in (A/B). Cytoplasmic (CYTO) and mitochondrial-enriched fractions (MITO) as indicated. HSP60 used as a mitochondrial loading control and HSP90 used as a cytoplasmic loading control. Protein MW ladder (in kilodaltons) as depicted. Representative of N=8 independent experiments. L. Quantification of TIGAR expression normalised to HSP60 in mitochondrial-enriched fractions from p53 WT and p53 KO HepG2 cells (G). N-numbers as in (G) and presented and analysed as in (B). ns- not significant, **p<0.01, ***p<0.001.

Functionally, although *p53* WT and *p53* KO HepG2 cells accumulated similar amounts of lipid after 48hrs of growth within HFHS medium, HepG2 *p53* KO cells exhibited significantly increased levels of ROS that were not observed within HFHS-treated HepG2 *p53* WT cells (Fig. 4E/F). This response was also observed after siRNA-mediated depletion of *TP53* in the HepG2 cells—although basal p53 protein levels were difficult to detect compared with downstream targets in the HepG2 cells, as has been previously reported^57^—or following siRNA-mediated depletion of *Trp53* within the Hep53.4 cells (Figs. 4G-J and S3A-E). Treatment with the anti-oxidant N-acetyl-cysteine (NAC) was sufficient to restore redox control to HFHS-treated *TP53* KO HepG2 cells without altering lipid accumulation, consistent with p53-mediated redox control supporting this process (Fig. S3F/G). Conversely, siRNA-mediated knockdown of *TIGAR/Tigar* in both *p53* WT HepG2 and Hep53.4 cells phenocopied CRISPR-mediated p53 depletion, resulting in significantly increased ROS in HFHS conditions without altering lipid accumulation (Fig. 4G-J).

TIGAR has been reported to exert antioxidant activities both in the cytoplasm as a consequence of its fructose-2,6-bisphosphatase activity and at the mitochondria independent of this function^47, 48^. In both HepG2 and Hep53.4 cells we observed a clear double band for TIGAR on western blots (Figs. 4A/C and S3A/D). Interestingly, these bands were found to preferentially localise to either cytoplasmic or mitochondrially-enriched fractions and the clearest TIGAR accumulation we observed was within the mitochondrial-enriched cell fraction in HepG2 cells treated with HFHS medium (Fig. 4K/L). Of note, these findings did not generalise to other *TP53* wildtype human cancer cell lines we examined, including the SK-Hep-1 liver cancer cell line or the colorectal carcinoma HCT116 cell line. The majority of TIGAR in these cells was cytoplasmic, the p53 pathway was not induced by HFHS diet treatment, and perturbing p53 with siRNA in SK-Hep-1 cells did not alter cellular ROS levels (Fig. S3H-J). Although derived from the liver of a patient with adenocarcinoma, SK-Hep-1 cells are endothelial in origin, with gene expression and morphological characteristics more consistent with liver sinusoidal endothelial cells (LSECs) rather than with an epithelial hepatocyte origin^58^. We believe this difference explains the observed lack of a p53-TIGAR response to HFHS conditions within SK-Hep-1 cells, as in the colorectal HCT116 cells, compared with the hepatocellular HepG2 and Hep53.4 cell lines.

Based on these findings, we concluded that p53-mediated induction of TIGAR represents an important aspect of redox control within the hepatocellular compartment during the response to HFHS conditions *in vitro*.

### Loss of TIGAR increases ROS and accelerates liver disease in leptin receptor-deficient (Lepr^db/db^) mice

Based on our *in vitro* data identifying TIGAR as an important mediator for managing the redox response to HFHS diet conditions, we sought to validate its importance in liver disease more broadly by returning to an *in vivo* model. Since the leptin-deficient genetic model of obesity yielded a rapid and potent example of repercussions for liver p53 loss (Fig. 3), we decided to examine the consequences of Tigar loss in a similar setting. Owing to the fact that *Tigar* and leptin (*Lep*) share murine chromosome 6, instead of attempting to create a *Tigar-*deficient*; Lep^ob^*compound mouse, we instead generated leptin receptor-deficient (*Lepr^db/db^*(*db/db*)) mice harbouring whole body deletion of *Tigar* (*Tigar^-/-^; db/db* mice). As with *ob/ob* mice, *db/db* mice develop progressive obesity due to hyperphagia alongside hyperglycaemia and frank diabetes^53^. The diabetic condition of *db/db* mice is generally more severe than that observed in *ob/ob* mice and lifespan is markedly decreased^53, 59^. This is especially true within male *db/db* mice, where mean survival has been reported to be only 14 months, compared with 19 months for female *db/db* mice^59^. Considering the already rapid lethality that we observed in *ob/ob; Albumin-Cre*; *Trp53^FL/FL^*mice (Fig. 3A), we decided to focus on liver disease development within cohorts of female *Tigar^-/-^; db/db* mice compared with *Tigar* WT; *db/db* control mice to ameliorate the potentially confounding effects of accelerated lethality expected in *db/db* male mice. Young (100-day-old) *Tigar^-/-^; db/db* female mice exhibited histopathological features of MASLD including steatosis and hepatocyte ballooning alongside significantly increased IHC staining for MDA—indicative of elevated lipid peroxidation (Fig. 5A/B). In addition, *Tigar^-/-^; db/db* female mice also had greater ALT and AST serum enzyme activity than *Tigar* WT; *db/db*, consistent with established liver damage in these mice (Fig. 5C/D). These findings are consistent with our data from *ob/ob; Albumin-Cre*; *Trp53^FL/FL^* mice, albeit from a cohort examined at a much earlier time point and with a differing genetic driver of obesity. In contrast with *ob/ob; Albumin-Cre*; *Trp53^FL/FL^* mice, 100-day-old *Tigar^-/-^; db/db* female mice did not exhibit increased levels of picrosirius red staining for fibrosis or have altered levels of lipogranulomas formation compared with age-matched *Tigar* WT; *db/db* mice (Fig. 5E/F). These differences could be a consequence of the earlier timepoint examined in *Db* mice (∼100 days vs ∼300 days), but could also be related to reported differences in the manifestation of MASLD/MASH between *ob/ob* and *db/db* mouse strains^60^.

**Figure 5:**
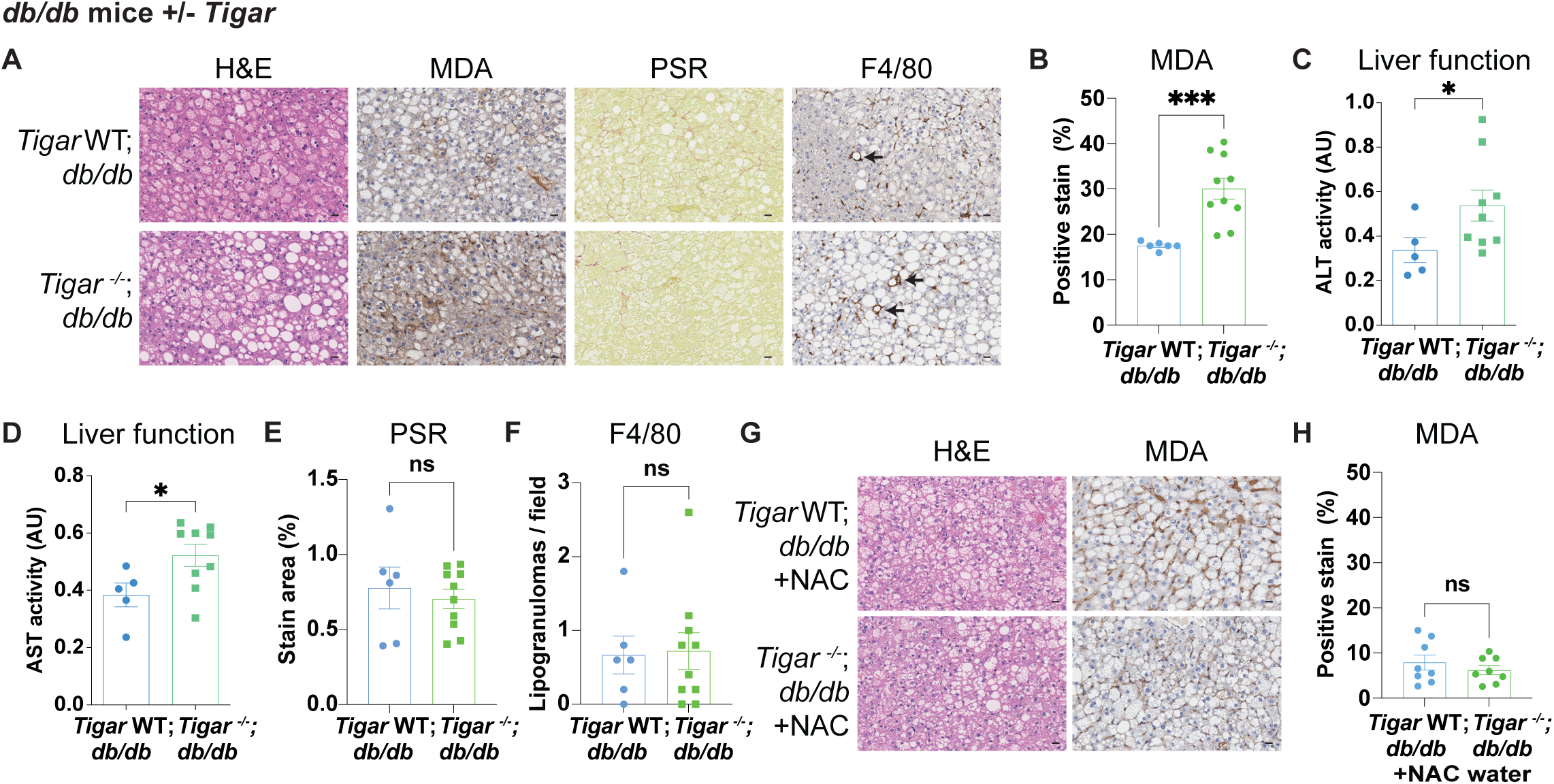
TIGAR limits ROS and protects liver function in leptin receptor-deficient (*Lepr^db/db^*) mice. A. Representative H&E images, IHC staining for malondialdehyde (MDA), staining for fibrosis (picrosirius red/PSR), and IHC staining for macrophages (F4/80) in female *Lepr^db/db^* (*Tigar* WT; *db/db*) and *Tigar^-/-^*; *Lepr^db/db^* (*Tigar*^-/-^; *db/db*) mice at 100 days old. Representative of N=6 *Tigar* WT; *db/db* and N=10 *Tigar*^-/-^; *db/db* mice. Macrophage-engulfed steatotic hepatocytes (lipogranulomas) depicted by arrows in F4/80 images. Scale bars 20 μm. B. Quantification of MDA to assess lipid peroxidation in *Lepr^db/db^*(*Tigar* WT; *db/db*) and *Tigar^-/-^*; *Lepr^db/db^* (*Tigar*^-/-^; *db/db*) mice at 100 days old as in (A). N-numbers as in (A). Data presented as mean +/- SEM and analysed using an unpaired t-test with Welch’s correction. **p<0.01. C/D. Plasma activity (mU/mL) of alanine (C) and aspartate (D) transaminase (ALT/AST) to assess liver function in female *Lepr^db/db^* (*Tigar* WT; *db/db*) and *Tigar^-/-^*; *Lepr^db/db^* (*Tigar*^-/-^; *db/db*) mice at 100 days old. N=5 *Tigar* WT; *db/db* and N=9 *Tigar*^-/-^; *db/db* mice. Each data point represents the mean from technical duplicates per mouse. Data obtained from a subset of mice in (A/B), with plasma not obtained from N=1 in each cohort. Data presented and analysed as in (B). *p<0.05. E/F. Quantification of PSR to assess fibrosis (E) and F4/80 for lipogranulomas (F) as in (B). Data presented and analysed as in (B). ns- not significant. G. Representative H&E images and IHC staining for malondialdehyde (MDA) in female *Lepr^db/db^* (*Tigar* WT; *db/db*) and *Tigar^-/-^*; *Lepr^db/db^* (*Tigar*^-/-^; *db/db*) mice at 100 days old and treated with N-Acetyl-Cysteine (NAC) in the drinking water from 42 days of age. Representative of N=7 *Tigar* WT; *db/db* + NAC and N=7 *Tigar*^-/-^; *db/db* + NAC mice. Scale bars 20 μm. H. Quantification of MDA to assess lipid peroxidation in *Lepr^db/db^*(*Tigar* WT; *db/db*) and *Tigar^-/-^*; *Lepr^db/db^* (*Tigar*^-/-^; *db/db*) mice treated with N-Acetyl-Cysteine (NAC) in the drinking water from 42 days of age (as in G). N=7 *Tigar* WT; *db/db* + NAC and N=7 *Tiga*r^-/-^; *db/db* mice + NAC. Data presented and analysed as in (B). ns- not significant.

To test whether redox control, as opposed to other functions of TIGAR, underpinned our findings *in vivo*, we supplemented the drinking water of *Tigar^-/-^; db/db* and *Tigar* WT; *db/db* female mice with the anti-oxidant NAC beginning at 6 weeks of age and assessed liver disease development within these cohorts when the mice reached 100 days of age. In NAC-treated mice, the histology of *Tigar^-/-^; db/db* and *Tigar* WT; *db/db* mice were similar and IHC staining for MDA was low for both, suggesting that NAC supplementation could prevent the increased lipid peroxidation and histopathological features of MASLD evident within *Tigar^-/-^; db/db* mice—at least at the timepoint examined (Fig. 5G/H). Staining for fibrosis was similarly low in both NAC-treated cohorts, although we did observe a small increase in the number of lipogranulomas per field within *Tigar^-/-^; db/db* mice treated with NAC compared with NAC-treated *Tigar* WT; *db/db* controls (Fig. S4A-C).

Together, these findings suggest that *Tigar* is important for limiting liver disease progression within the *db/db* genetic model and that targeting ROS directly with an antioxidant can alleviate detrimental effects of *Tigar* loss *in vivo*.

## DISCUSSION

During the transition from MASLD to MASH, the liver undergoes successive alterations that are reminiscent of key steps of tumourigenesis. Progressive rewiring of hepatic metabolism, significant inflammation, development of tissue damage, and engagement of cancer-associated signalling programmes all contribute to the multi-faceted disease state that defines advanced MASH^8, 9^. Many of these attributes are also important hallmarks of cancer^11^. With this overlap in mind, we have interrogated the activity of the tumour suppressor protein p53 throughout MASLD and MASH development. Our findings identify p53-mediated redox control and the induction of TIGAR as important protective features of the hepatic p53 response both *in vivo* and in human and murine HCC cell lines *in vitro*.

We report that p53 activation is an early feature of MASLD pathogenesis that increases alongside disease progression in p53 reporter mice. Across experimental models, we have found that loss of hepatic p53 is associated with worsening liver disease that is characterised by liver damage, chronic inflammation, fibrosis, and high levels of ROS. In *Trp53/TP53* wildtype murine and human HCC cell lines, we show that the antioxidant gene *Tigar/TIGAR* is induced in a p53-dependent manner after HFHS treatment, localises to the mitochondria, and supports redox control. In response to HFHS medium, loss of either p53 or TIGAR exacerbates redox stress and this can be rescued with N-acetyl-cysteine supplementation, confirming the role of TIGAR in supporting p53-mediated ROS control in this setting. Consistent with these observations, in the *db/db* genetic model of obesity, we show that *Tigar*-deficient mice exhibit evidence of enhanced liver disease, including increased ROS and liver damage from an early age that can also be rescued by N-acetyl-cysteine supplementation. Our findings support the conclusion that TIGAR-mediated redox control, likely alongside additional functions of p53, protects the liver against deleterious consequences of chronic consumption of a high fat and high sugar ‘Western’ diet. This protection is lost in the absence of liver p53 or TIGAR but can be partially restored with extended antioxidant treatment.

Our results add nuance to reports in mouse models and human patients where p53 induction and/or increased p53 pathway activity are associated with MASLD to MASH progression^20, 21, 36, 61^. We also observe greater p53 activity in advanced vs early stages of diet-induced liver disease. Nevertheless, induction of p53 in our reporter mice is clearly a much earlier feature of adaptation to HFHS diet stress, at least in male mice, and occurs prior to evidence of, for example, significant glucose intolerance that would indicate advanced metabolic syndrome, T2DM, or the onset of MASH. Whether this is also true within female mice is not yet clear, and it would be interesting to examine whether noted sex differences in the p53 response that we observed reflect different underlying biology in future studies. Nevertheless, combined with earlier reports, our findings reported here suggest a dynamic interaction may exist between diet-induced stress and p53 activity. In this model, the modest activation of p53 we observe early in MASLD disease aetiology may be sufficient to coordinate redox protection, but this could eventually switch into a pro-disease progression paradigm later when p53 signalling increases above a critical threshold and p53-mediated apoptosis, for example, is achieved^36^. Such a model would be consistent with more general literature on the threshold mechanisms inherent in determining the p53 response^62^.

Focusing on this point further, it is tempting to propose metabolic zonation as a potential mediator of the p53 response during MASLD to MASH progression. Diet-induced steatosis begins pericentrally, as we observed within our HFHS diet model and as is reported in human patients^63^. This region of the liver maintains low oxygen tension compared with periportal regions of the liver^63^. Even mild hypoxia is known to alter p53 activity, including changing the regulation of p53-mediated apoptosis and altering the p53 response to ROS^64, 65^. One of the main features of MASLD progression to MASH is the overall expansion of steatosis from a zonally-restricted pericentral localisation to a global distribution, with lipid accumulation, inflammation, and fibrosis occurring across the liver lobule, including within more oxygenated areas of the liver where p53 activity is normalised. Future work could examine this possibility, for example by mapping the spatio-temporal distribution of p53 activation throughout MASLD progression to MASH. This would provide insight into the molecular mechanisms that underpin p53 activity during liver disease.

Taken together, our results underscore the importance of p53 and TIGAR for protecting the liver against damage associated with diet-induced liver disease. They also suggest that antioxidant interventions may have efficacy in preventing some of the deleterious aspects of MASH. Interestingly, considering that metabolic syndrome is a multi-organ condition, our findings also suggest that nuances of the p53 response to diet stress are disease-state specific and potentially not shared across all metabolic organs.

## MATERIALS AND METHODS

### Mice

Procedures involving mice were performed under Home Office licence numbers 70/8645, 70/8468, PP6345023 and PP1389725 as previously described^50^. Experiments were conducted in accordance with the Animals (Scientific Procedures) Act 1986 and the EU Directive 2010 and were sanctioned by Local Ethical Review Process (University of Glasgow) and The Francis Crick Institute. Mice were housed on a 12-hour light/12-hour dark cycle in groups of 3-5 as much as possible. Mice were provided with environmental enrichment in the form of Sizzle-Nest bedding and polycarbonate tunnels, normal chow diet, and water ad libitum. Non-aversive handling techniques were utilised throughout each experiment. Mice were genotyped by Transnetyx (Cordova, TN).

*p53^FL/FL^* (*Trp53*^tm1Brn^), *Albumin*-Cre (Speer6-ps1^Tg(Alb-cre)21Mgn^), PG13-iRFP (*Hprt^Tm1(p53RE-iRFP [pg13])Bea^*), and *Tigar^-/-^*mice were described previously ^49, 50, 66, 67^ and were fully backcrossed to C57BL6/J (N10). *Lep^ob^(B6.Cg-Lep^ob^/J)* and *Lepr^db^* (BKS.Cg- *Dock7^m^*+/+*Lepr^db^*J) heterozygous mice were purchased from Charles River (Strain codes: 606 and 607). *Albumin*-Cre; *Trp53^FL/FL^*compound mice were previously described^43^. *Lep^ob/ob^*; *Albumin*-Cre; *Trp53^FL/FL^* mice were generated by interbreeding *Lep^ob/WT^* and *Albumin*-Cre; *Trp53^FL/FL^* mice. *Lepr^db/db^*; *Tigar^-/-^* mice were generated by interbreeding *Lepr^db/WT^* and *Tigar^WT^*^/-^ mice.

Within experimental cohorts, mice were age and littermate-matched as much as possible, and all were started on experimental procedures and/or imaging prior to 6 months of age. In the imaging studies, a subset of the mice were sampled at either 100 days (N=5 male chow fed mice, N=3 male HFHS fed mice, N=3 chow fed female mice, or N=2 HFHS fed female mice) or 150 days into the experiment (N=3 HFHS fed male mice). In addition, two female mice on the HFHS diet developed skin conditions that required them to be euthanised. These animals were included until the point of euthanasia, and removed from the longitudinal data after these instances.

The presented weight measurements in Figure 1 were obtained from the reported imaging cohorts, but timepoints prior to 150 days reflect only a subset of these mice, as these data were not initially collected from the first groups of experimental animals.

Imaging cohorts (Fig. 1) included both male and female mice. These were either hemizygous (males) or homozygous (females) for the PG13-iRFP allele. Based on imaging results that indicated a preponderance of p53 activity in males during the response to HFHS diet, the functional consequences of liver *Trp53* deletion were examined in only male cohorts (Fig. 2). Experiments involving *Lep^ob^* mice (Fig. 3) were undertaken in both male and female mice. Considering the rapid sex-independent lethality observed within *Lep^ob/ob^*; *Albumin*-Cre; *Trp53*^FL/FL^ mice, experiments examining *Tigar*-deficiency within the more aggressive *Lepr^db^* model were undertaken in only females, where the normal disease latency is significantly longer^59^.

Downstream analyses were performed on a random order of samples by researchers blinded to the genotype and treatment regime of a given sample until the summation of results.

### Long-term diet experiments

For long-term diet experiments, male 65–75 day old *Trp53* wildtype; *Albumin*-Cre and *Trp53*^FL/FL^; *Albumin*-Cre mice were shifted onto either an obesogenic high-fat and high-sugar (HFHS) diet or left on normal mouse chow (control diet). HFHS diet (AIN-76A, 58R3) was purchased from TestDiet and contained 59% energy from fat, 15% energy from protein, 25% energy from carbohydrates, was rich in sucrose, and did not contain cholesterol. The control normal mouse chow (DS801752G10R, Special Diet Services/SDS) contained 9% energy from fat, 21% energy from protein, 70% energy from carbohydrates and reduced sucrose.

For imaging long-term diet experiments, male and female 65–75-day old p53 reporter (PG13-iRFP) mice were all shifted onto a purified control maintenance diet suitable for imaging mice (TestDiet, AIN-93M, 58M1) rather than normal mouse chow. This diet change was done at least 7 days prior to imaging mice in order to eliminate non-specific fluorescence in the gut prior to obtaining baseline p53 levels. Mice in imaging experiments continued to be maintained on either control imaging diet or were shifted onto the HFHS diet for the duration of the experiment thereafter. As with the purified control maintenance diet, the HFHS diet was also confirmed to not cause non-specific fluorescence in the gut.

### In vivo imaging

Mice were imaged longitudinally on a Pearl Impulse Small Animal Imaging System (LI-COR) as previously described^68^. For quantification, the PG13-iRFP signal registered within the liver region of each mouse was normalised to the individual initial baseline liver-region signal for each mouse, taken at the start of the experiment. For a small number of experimental mice, the observed iRFP signal of the initial baseline scan was either transiently very low (N=3/42) or very high (N=1/42) for one imaging session compared with subsequent scans. In these cases, the liver region signal observed one week later was instead used to define the baseline for longitudinal analysis. Scan images are presented with false colour LUTs. Imaging parameters were kept constant across time points and mice within an experiment. For quantification, Image Studio software (LI-COR, V5.5) was used.

### *Ex vivo* imaging of tissues

Harvested tissues were fixed overnight in 10% neutral buffered formalin (Solmedia), assembled onto a 10 cm petri dish, and imaged on a Pearl Impulse Small Animal Imaging System (LI-COR) as for *in vivo* imaging described above. Imaging parameters were kept constant across samples and scans were analysed using Image Studio software (LI-COR, V5.5). Scan images are presented with false colour LUTs which were held constant across all images shown.

### Oral Glucose Tolerance Test

Prior to the oral glucose tolerance test, mice were fasted for a period of 4 hours. Sterile glucose (60% w/v in water) was administered in a bolus of 3 g/kg via oral gavage per mouse, based on weights obtained at the start of the fasting period. Blood glucose levels were measured with an Accu-Chek blood glucose meter (Accu-Chek UK) using blood obtained from the tail vein before oral gavage (time 0), and at 15 mins, 30 mins, 45 mins, 60 mins, and 120 mins after administration of the glucose bolus.

### Liver function tests

Alanine transaminase (ALT) and aspartate aminotransferase (AST) activity were measured in EDTA-treated plasma using kits from Abcam (ab105134 and ab138878) according to the manufacturer’s instructions. Samples were run together, analysed in duplicate wells per mouse, and the mean value of these technical replicates was used for subsequent analysis.

### Histology and Immunohistochemistry (IHC)

All haematoxylin & eosin (H&E), immunohistochemistry (IHC), and picrosirius red (PSR) staining took place on 4 µm formalin fixed paraffin embedded sections (FFPE), which had previously been heated in an oven at 60⁰C for 2 hours. Oil-red-o (ORO) staining was performed on 10 μm frozen sections that were first fixed for 5 minutes in 10% neutral buffered formalin (Solmedia), rinsed in tap water, and then briefly rinsed in 60% isopropanol (Fisher Chemicals).

The following antibodies were stained on a Leica Bond Rx autostainer: F4/80 (ab6640, Abcam) and malondialdehyde (MDA) (ab243066, Abcam). All FFPE sections underwent on-board dewaxing (AR9222, Leica) and epitope retrieval using appropriate retrieval solutions. Sections for F4/80 staining underwent epitope retrieval using enzyme 1 solution (AR9551, Leica) for 10 minutes at 37°C. Sections for MDA staining underwent epitope retrieval with ER2 solution (AR9640, Leica) for 20 minutes at 95°C. Sections were washed with Leica wash buffer (AR9590, Leica) before peroxidase block was performed using an Intense R kit (DS9263, Leica). MDA sections had an additional Mouse Ig blocking step (MKB-2213, Vector Labs) applied for 20 minutes. All sections were then rinsed with wash buffer before primary antibody application at the optimal dilution (F4/80: 1/200; MDA: 1/250). The sections were rinsed with wash buffer and appropriate secondary antibody applied for 30 minutes. MDA sections had Mouse Envision (K4001, Agilent) applied and sections for F4/80 had Rat ImmPRESS secondary solution (MP-7404, Vector Labs) applied. Sections were rinsed with wash buffer before being visualised using DAB and counterstained with haematoxylin in the Intense R kit.

H&E staining was performed on a Leica autostainer (ST5020). Sections were dewaxed in xylene, taken through graded ethanol solutions and stained with Haem Z (RBA-4201-00A, CellPath) for 13 mins. Sections were washed in water, differentiated in 1% acid alcohol, and washed. Nuclei were blued using Scott’s Tap Water Substitute (in-house). After washing with tap water, sections were placed in Putt’s Eosin (in-house) for 3 minutes to complete the staining.

Staining for PSR and ORO was performed as previously described^43^.

All stained sections except for those from experiments involving *Lep^ob^* mice (Fig. 3) were scanned at 20x magnification using a Leica Aperio AT2 slide scanner (Leica Microsystems, UK) prior to subsequent analysis of scanned slides as described below.

### IHC and Staining Quantification

All quantification was undertaken on a random order of samples blinded to the genotype and treatment regime of a given sample until the summation of results.

Quantification for MDA, F4/80-positive lipogranulomas and PSR staining in Figs. 1, 2, and 5 were carried out using QuPath software (v0.5.1)^69^. The same parameters per stain were used for all samples. For PSR and MDA staining, random 20x magnification images were selected taking care to avoid any vessels where these stains were excessively high. The average percentage of positively stained area was calculated for each mouse. F4/80 staining attributable to hepatocyte-engulfing macrophages was scored by hand from five random 20x magnification images per slide in QuPath and averaged per mouse sample.

Quantifications in Fig. 3 were performed as previously described^43^.

### Cell culture

HepG2 (HB-8065), SK-Hep-1 (HTB-52), and HCT116 (CCL-247) cells were obtained from ATCC but were not authenticated for this study. Hep53.4 cells (Cellosaurus CVCL_5765) were obtained from Cytion (Product number 400200) but were not authenticated for this study. HepG2 *TP53* knockout (p53 KO) cells were generated through CRISPR-mediated targeting of *TP53* as previously described^70^. Mycoplasma testing was performed on all cells when they were thawed and semi-regularly thereafter using the MycoAlert PLUS Mycoplasma Detection Kit (Lonza LT07-703). Independent experiments were performed on cells treated with siRNA and/or compounds from separate passages of each cell line, with technical replicates used to confirm the effect of treatment conditions and/or siRNA-mediated gene knockdown. Stock flasks were maintained in DMEM, low glucose, pyruvate, no glutamine, no phenol red (Gibco cat# 11880036) supplemented with 2 mM L-Glutamine (Gibco cat# 25030032), penicillin/streptomycin (Gibco cat# 15070063), and 5% FBS (Merck cat# F7524). Cells were cultured at 37°C in a humidified atmosphere of 5% CO_2_.

### Transfection with siRNA

Studies utilising siRNA knockdown were performed as previously described^70^ using ON-TARGETplus SMARTpool constructs (Horizon Discovery Biosciences) including the non-targeting siRNA control pool (D-001810-10) and constructs targeting human *TP53* (L-003329-00), human *TIGAR* (L-020597-01), mouse *Trp53* (L-040642-00), and mouse *Tigar* (L-056801-01). Cell lines were transfected with constructs at a final concentration of 20 nM using DharmaFECT 4 transfection reagent (T-2004) and the manufacturer’s recommended reverse transfection procedure (Horizon Discovery Biosciences).

### *In vitro* HFHS medium formulation

To create the HFHS medium formulation, a concentrated lipid mixture stock solution of fatty acids conjugated to fatty acid-free BSA (10% solution in PBS) (Merck cat# A1595) was created as previously described^71^. The stock solution was added at 1% v/v to Plasmax cell culture medium (CancerTools cat# 156371) supplemented with 5% FBS (Merck cat# F7524) and penicillin/streptomycin (Gibco cat# 15070063). The resulting high fat medium was further supplemented with an additional 20 mM D-Glucose (Merck cat# G7528) and 25mM D-Fructose (Thermo Scientific cat# A17718.30) to provide the high sugar component of the HFHS medium. The final concentrations of fatty acids used in the HFHS medium were: 150 µM palmitate (Merck cat# P5585), 150 µM stearic acid (Thermo Scientific cat# A12244.06), 25 µM linoleic acid (Merck cat# L9530), 2.8 µM linolenic acid (Thermo Scientific cat# 215040050), 0.14 µM oleic acid (Thermo Scientific cat# 270290050) and 2 µM arachidonic acid (Sigma, cat# 10931). These amounts were chosen to broadly match the lipid distribution found in the murine HFHS diet. For control-treated cells, Plasmax supplemented with 1% v/v of the 10% fatty acid-free BSA solution in PBS (Merck cat# A1595) supplemented with 10% Ethanol (Fisher Scientific cat# 10680993) (0.1% v/v final concentration) was used to mirror the ethanol used to dissolve fatty acids for BSA conjugation in HFHS medium formulation.

For experiments utilising HFHS medium, cells were plated at an assay-dependent dilution in the maintenance DMEM medium as described above. For flow cytometry and western blotting, 1.25*10^5^ cells/well were seeded in 6-well plates (Thermo Scientific cat# 140675) and for mitochondrial enrichment, 3.0*10^6^ cells were seeded in 150 mm dishes (Thermo Scientific cat# 168381). 24–48 hours after seeding, cells were treated with either HFHS Plasmax or Plasmax supplemented with BSA (control) and left for a further period of 48 hours in treatment medium prior to analysis via Flow Cytometry, Western Blotting, or mitochondrial enrichment. In cells transfected with siRNA, reverse transfection was undertaken when cells were seeded in 6-well plates, and the transfection mixture was left on the cells until treatment medium was added for each condition.

### Antioxidant treatment with N-acetyl-cysteine

For experiments involving the antioxidant N-Acetyl-cysteine (NAC), cells were treated with 2 mM NAC (Sigma cat# A7250) dissolved in PBS for a period of 48 hours. NAC treatment was added directly to the HFHS medium used for treated samples.

### Flow Cytometry

Adherent cells were detached with Trypsin-EDTA solution (Merck cat# T4174), dissociated into single cells, and resuspended in phenol red-free DMEM (Gibco cat# 11880036). Cells were labelled with CellROX Deep Red reagent (7.5 μM, Thermo Fisher Scientific cat# C10422) and BODIPY 493/503 (5 μM, Thermo Fisher Scientific cat# D3922) for 10 minutes at room temperature in a 1:1 mixture of phenol red-free DMEM and a staining solution containing the dyes in PBS without calcium and magnesium (Corning cat# 21-040-CV) supplemented with 2.5% w/v BSA (Merck cat# 810533). 4′,6-Diamidino-2-phenylindole dihydrochloride (DAPI, Merck cat# D9542) was added to a final concentration of 1 μg/ml to each sample and was used to identify viable cells for analysis. Single cells were analysed on a MACSQuant Analyzer 10 (Miltenyi Biotec) using unstained and hydrogen peroxide-treated cells as controls. At least 10,000 events were collected for each sample. Data were analysed using FlowJo 10.10.0 (BD Biosciences). Unless otherwise stated, median fluorescence intensity values were obtained and compared across samples.

### Mitochondrial Enrichment

The isolation of cytoplasmic and mitochondrial-enriched fractions from cells was performed as described previously^72^. Briefly, cells were scraped in isotonic HIM buffer (200 mM mannitol, 1 mM EGTA, 70 mM sucrose, 10 mM HEPES), passed through a cell homogeniser with a 16 µm bead and pelleted by centrifugation at various speeds to collect individual enriched fractions. The cytoplasmic fraction was retain in HIM buffer and the mitochondrial-enriched pellet was resuspended in NP-40 lysis buffer (Fisher Scientific cat# 15403489) supplemented with PhosSTOP (Merck cat# 4906845001) and the cOmplete Ultra EDTA-free protease inhibitor cocktail (Merck cat# 05892791001).

### Western Blotting

Western blotting was performed as previously described^70^ with the following modifications:

Protein lysates were prepared using NP-40 lysis buffer as in mitochondrial enrichment experiments. Protein concentration was determined in each sample using a Pierce BCA Protein Assay kit (cat# 23227), equivalent concentrations were loaded in each well, and total protein was confirmed following transfer using a Pierce Reversible Protein Stain Kit (Fisher cat# PI24580). Blocking was performed in a TBS solution containing 5% BSA (Merck cat# 810533) and 0.1% Tween-20 (Merck cat# P1379), and wash steps were performed in TBS with 0.1% Tween-20 (TBS-T).

Primary and Secondary antibodies were used as indicated in the tables below.

Proteins were detected using either an Odyssey DLx or Odyssey XF Imaging System (LI-COR Biosciences) and analysed using ImageStudio 5.5 software (LI-COR Biosciences).

### Primary antibodies

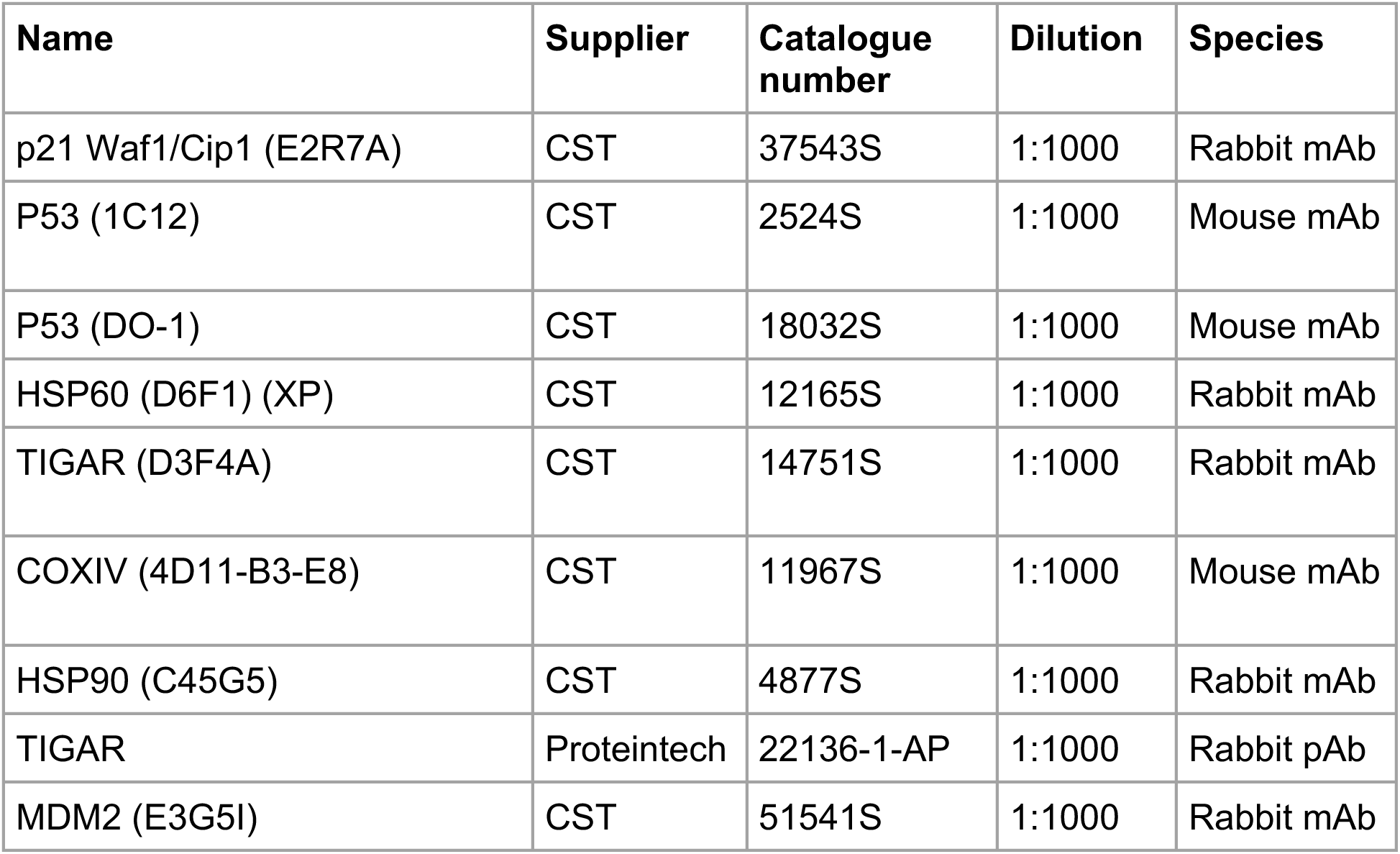

### Secondary antibodies

| Name | Supplier | Catalogue number | Dilution |
| --- | --- | --- | --- |
| IRDye 680LT Donkey anti-Mouse | Li-COR | 926-68022 | 1:20,000 |
| IRDye 800CW Donkey anti-Rabbit | Li-COR | 926-32213 | 1:20,000 |

### Quantitative RT-PCR

Quantitative RT-PCR analysis of HepG2 cells was undertaken as previously described^43^, using Taqman Fast Advanced Master Mix and the Taqman Gene Expression Assays indicated in the table below.

| Gene | Supplier | Assay ID |
| --- | --- | --- |
| *ACTB* | Fisher Scientific | Hs01060665_g1 |
| *TP53* | Fisher Scientific | Hs01034249_m1 |
| *CDKN1A* | Fisher Scientific | Hs00355782_m1 |
| *TIGAR* | Fisher Scientific | Hs00608644_m1 |

### Data plotting and statistical analysis

Data were plotted using GraphPad Prism 10 (GraphPad). The statistical analysis for each experiment was performed using the test indicated in the relevant figure legend and multiplicity-adjusted p-values using the built-in analysis tools of Prism 10. Statistical tests were chosen based on standard tests utilised in the field and the nature of the comparison being made. Underlying assumptions for these tests were assumed to be met although not explicitly examined. Figures were prepared using Adobe Illustrator 2024 (Adobe). Data are presented as mean with min/max floating bars and individual data points (Figs. 1 and S1) and mean +/- SEM with individual data points (Fig. 2 onwards). P-values are denoted as follows: ns- not significant, *p<0.05, **p<0.01, ***p<0.001, ****p<0.0001.

## ACKNOWLEDGEMENTS

We thank the Core Facilities and Advanced Technologies at the CRUK Scotland Institute, and in particular the Biological Services facilities staff and the histology team. We also thank the Crick BRF for helping with animal experiments, and the Crick EHP for histopathology support. We thank Saverio Tardito (CRUK Scotland Institute) for fruitful discussions about the project and the initial provision of Plasmax medium for our studies, Thomas G. Bird (CRUK Scotland Institute / University of Edinburgh) for providing Hep53.4 cells and for thoughtful comments on the manuscript, Catherine Winchester (CRUK Scotland Institute) for insightful comments on the manuscript, and Mark Thomas Shaw Williams (Glasgow Caledonian University) for constructive feedback on the project and helpful comments on the manuscript.

## AUTHOR CONTRIBUTIONS

CRediT (Contributor Roles Taxonomy) author statement:

**CIW:** Methodology, Investigation, Formal Analysis, Writing-Original Draft Preparation; **ECC:** Methodology, Investigation, Formal Analysis, Writing-Review & Editing; **DA, NC, LB, MH, VM, DMW:** Investigation, Writing- Review & Editing; **KB, KHV:** Conceptualization, Supervision, Funding acquisition, Writing- Review & Editing; **TJH:** Conceptualization, Investigation, Formal Analysis, Writing- Original Draft, Supervision, Project Administration, Funding Acquisition.

## DATA AND MATERIALS AVAILABILITY

All data needed to evaluate the conclusions in the paper are present either within the paper itself or the included Supplementary Materials. Underlying raw data from this study are available from the corresponding author upon reasonable request.

## SUPPLEMENTAL MATERIALS

**Figures S1-S4**

**S1:** Liver p53 is engaged more strongly in male mice in response to a high fat and high sugar diet

**S2:** Liver p53 limits liver damage arising from a high fat and high sugar diet *in vivo*

**S3:** p53 supports redox control and engages TIGAR in *p53* WT HepG2 and Hep53.4 HCC cell lines in response to overfeeding *in vitro*

**S4:** N-acetyl-cysteine supplementation reduces ROS in leptin receptor-deficient *Lepr^db/db^* mice that lack *Tigar* expression

## REFERENCES

1. Burton, A. et al. Primary liver cancer in the UK: Incidence, incidence-based mortality, and survival by subtype, sex, and nation. JHEP Reports 3, 100232 (2021).

2. Huang, D. Q. et al. Changing global epidemiology of liver cancer from 2010 to 2019: NASH is the fastest growing cause of liver cancer. Cell Metab 34, 969–977.e2 (2022).

3. Rinella, M. E. et al. A multisociety Delphi consensus statement on new fatty liver disease nomenclature. Hepatology 78, 1966–1986 (2023).

4. Hester, D. et al. Among Medicare Patients With Hepatocellular Carcinoma, Non–alcoholic Fatty Liver Disease is the Most Common Etiology and Cause of Mortality. J Clin Gastroenterol 54, 459–467 (2020).

5. Asrani, S. K., Devarbhavi, H., Eaton, J. & Kamath, P. S. Burden of liver diseases in the world. J Hepatol 70, 151–171 (2019).

6. Caldwell, S. H., Crespo, D. M., Kang, H. S. & Al-Osaimi, A. M. S. Obesity and hepatocellular carcinoma. Gastroenterology 127, S97–S103 (2004).

7. Huang, P. L. A comprehensive definition for metabolic syndrome. Dis Model Mech 2, 231– 237 (2009).

8. Powell, E. E., Wong, V. W.-S. & Rinella, M. Non-alcoholic fatty liver disease. The Lancet 397, 2212–2224 (2021).

9. Loomba, R., Friedman, S. L. & Shulman, G. I. Mechanisms and disease consequences of nonalcoholic fatty liver disease. Cell 184, 2537–2564 (2021).

10. Younossi, Z. M. et al. The global epidemiology of NAFLD and NASH in patients with type 2 diabetes: A systematic review and meta-analysis. J Hepatol 71, 793–801 (2019).

11. Hanahan, D. & Weinberg, R. a. Hallmarks of cancer: the next generation. Cell 144, 646–74 (2011).

12. Brown, G. T. & Kleiner, D. E. Histopathology of nonalcoholic fatty liver disease and nonalcoholic steatohepatitis. Metabolism 65, 1080–1086 (2016).

13. Seki, S. et al. In situ detection of lipid peroxidation and oxidative DNA damage in non-alcoholic fatty liver diseases. J Hepatol 37, 56–62 (2002).

14. Xiong, X. et al. Mapping the molecular signatures of diet-induced NASH and its regulation by the hepatokine Tsukushi. Mol Metab 20, 128–137 (2019).

15. Torkamani, A., Topol, E. J. & Schork, N. J. Pathway analysis of seven common diseases assessed by genome-wide association. Genomics 92, 265–272 (2008).

16. Hoang, S. A. et al. Gene Expression Predicts Histological Severity and Reveals Distinct Molecular Profiles of Nonalcoholic Fatty Liver Disease. Sci Rep 9, 12541 (2019).

17. Kleinstein, S. E. et al. Whole-Exome Sequencing Study of Extreme Phenotypes of NAFLD. Hepatol Commun 2, 1021–1029 (2018).

18. Seillier, M. et al. Defects in mitophagy promote redox-driven metabolic syndrome in the absence of TP53INP1. EMBO Mol Med 7, 802–818 (2015).

19. Setoyama, D., Fujimura, Y. & Miura, D. Metabolomics reveals that carnitine palmitoyltransferase-1 is a novel target for oxidative inactivation in human cells. Genes to Cells 18, 1107–1119 (2013).

20. Tomita, K. et al. p53/p66Shc-mediated signaling contributes to the progression of non-alcoholic steatohepatitis in humans and mice. J Hepatol 57, 837–843 (2012).

21. Aravinthan, A. et al. Hepatocyte senescence predicts progression in non-alcohol-related fatty liver disease. J Hepatol 58, 549–556 (2013).

22. Kung, C.-P. et al. The P72R Polymorphism of p53 Predisposes to Obesity and Metabolic Dysfunction. Cell Rep 14, 2413–2425 (2016).

23. Bonfigli, A. R. et al. The p53 codon 72 (Arg72Pro) polymorphism is associated with the degree of insulin resistance in type 2 diabetic subjects: a cross-sectional study. Acta Diabetol 50, 429–436 (2013).

24. Boutelle, A. M. & Attardi, L. D. p53 and Tumor Suppression: It Takes a Network. Trends in Cell Biology vol. 31 298–310 Preprint at 10.1016/j.tcb.2020.12.011 (2021).

25. Eriksson, S. E., Ceder, S., Bykov, V. J. N. & Wiman, K. G. p53 as a hub in cellular redox regulation and therapeutic target in cancer. J Mol Cell Biol 11, 330–341 (2019).

26. Berkers, C. R., Maddocks, O. D. K., Cheung, E. C., Mor, I. & Vousden, K. H. Metabolic regulation by p53 family members. Cell Metab 18, 617–633 (2013).

27. Levine, A. J. p53: 800 million years of evolution and 40 years of discovery. Nat Rev Cancer 20, 471–480 (2020).

28. Kastenhuber, E. R. & Lowe, S. W. Putting p53 in Context. Cell 170, 1062–1078 (2017).

29. Lu, W.-Y. et al. Hepatic progenitor cells of biliary origin with liver repopulation capacity. Nat Cell Biol 17, 971–983 (2015).

30. Katz, S. et al. Disruption of Trp53 in Livers of Mice Induces Formation of Carcinomas With Bilineal Differentiation. Gastroenterology 142, 1229–1239.e3 (2012).

31. Ferreira, D. M. S. et al. c-Jun N-Terminal Kinase 1/c-Jun Activation of the p53/MicroRNA 34a/Sirtuin 1 Pathway Contributes to Apoptosis Induced by Deoxycholic Acid in Rat Liver. Mol Cell Biol 34, 1100–1120 (2014).

32. Tian, X.-F., Ji, F.-J., Zang, H.-L. & Cao, H. Activation of the miR-34a/SIRT1/p53 Signaling Pathway Contributes to the Progress of Liver Fibrosis via Inducing Apoptosis in Hepatocytes but Not in HSCs. PLoS One 11, e0158657 (2016).

33. Bird, T. G. et al. TGFβ inhibition restores a regenerative response in acute liver injury by suppressing paracrine senescence. Sci Transl Med 10, eaan1230 (2018).

34. Buitrago-Molina, L. E. et al. The degree of liver injury determines the role of p21 in liver regeneration and hepatocarcinogenesis in mice. Hepatology 58, 1143–1152 (2013).

35. Marhenke, S. et al. p21 promotes sustained liver regeneration and hepatocarcinogenesis in chronic cholestatic liver injury. Gut 63, 1501–1512 (2014).

36. Farrell, G. C. et al. Apoptosis in experimental NASH is associated with p53 activation and TRAIL receptor expression. Journal of Gastroenterology and Hepatology (Australia) 24, 443– 452 (2009).

37. Kurinna, S. et al. p53 regulates a mitotic transcription program and determines ploidy in normal mouse liver. Hepatology 57, 2004–2013 (2013).

38. Huo, Y. et al. Protective role of p53 in acetaminophen hepatotoxicity. Free Radic Biol Med 106, 111–117 (2017).

39. Borude, P. et al. Pleiotropic Role of p53 in Injury and Liver Regeneration after Acetaminophen Overdose. American Journal of Pathology 188, 1406–1418 (2018).

40. Krizhanovsky, V. et al. Senescence of Activated Stellate Cells Limits Liver Fibrosis. Cell 134, 657–667 (2008).

41. Borude, P. et al. Pleiotropic Role of p53 in Injury and Liver Regeneration after Acetaminophen Overdose. Am J Pathol 188, 1406–1418 (2018).

42. Yu, G. et al. Loss of p53 Sensitizes Cells to Palmitic Acid-Induced Apoptosis by Reactive Oxygen Species Accumulation. Int J Mol Sci 20, 6268 (2019).

43. Humpton, T. J. et al. p53-mediated redox control promotes liver regeneration and maintains liver function in response to CCl4. Cell Death Differ 29, 514–526 (2022).

44. Lee, P., Vousden, K. H. & Cheung, E. C. TIGAR, TIGAR, burning bright. Cancer Metab 2, 1 (2014).

45. Bensaad, K. et al. TIGAR, a p53-inducible regulator of glycolysis and apoptosis. Cell 126, 107–20 (2006).

46. Anastasiou, D. et al. Inhibition of pyruvate kinase M2 by reactive oxygen species contributes to cellular antioxidant responses. Science 334, 1278–83 (2011).

47. Cheung, E. C., Ludwig, R. L. & Vousden, K. H. Mitochondrial localization of TIGAR under hypoxia stimulates HK2 and lowers ROS and cell death. Proc Natl Acad Sci U S A 109, 20491– 6 (2012).

48. Wang, X. et al. TIGAR reduces neuronal ferroptosis by inhibiting succinate dehydrogenase activity in cerebral ischemia. Free Radic Biol Med 216, 89–105 (2024).

49. Cheung, E. C. et al. TIGAR is required for efficient intestinal regeneration and tumorigenesis. Dev Cell 25, 463–77 (2013).

50. Humpton, T. J. et al. A noninvasive iRFP713 p53 reporter reveals dynamic p53 activity in response to irradiation and liver regeneration in vivo. Sci Signal 15, (2022).

51. Kheder, R. et al. Vitamin D3supplementation of a high fat high sugar diet ameliorates prediabetic phenotype in female LDLR–/–and LDLR+/+mice. Immun Inflamm Dis 5, 151–154 (2017).

52. Sutter, A. G., Palanisamy, A. P., Lench, J. H., Jessmore, A. P. & Chavin, K. D. Development of steatohepatitis in Ob/Ob mice is dependent on Toll-like receptor 4. Ann Hepatol 14, 735–43 (2015).

53. Coleman, D. L. Obese and diabetes: Two mutant genes causing diabetes-obesity syndromes in mice. Diabetologia 14, 141–148 (1978).

54. Vande Voorde, J., et al. Improving the metabolic fidelity of cancer models with a physiological cell culture medium. Sci Adv 5, eaau7314 (2019).

55. Kress, S. et al. p53 mutations are absent from carcinogen-induced mouse liver tumors but occur in cell lines established from these tumors. Mol Carcinog 6, 148–158 (1992).

56. Fischer, M. Census and evaluation of p53 target genes. Oncogene 36, 3943–3956 (2017).

57. Bai, J. & Cederbaum, A. I. Catalase Protects HepG2 Cells from Apoptosis Induced by DNA-damaging Agents by Accelerating the Degradation of p53. Journal of Biological Chemistry 278, 4660–4667 (2003).

58. Tai, Y. et al. SK-Hep1: not hepatocellular carcinoma cells but a cell model for liver sinusoidal endothelial cells. Int J Clin Exp Pathol 11, 2931–2938 (2018).

59. Sataranatarajan, K. et al. Rapamycin Increases Mortality in db/db Mice, a Mouse Model of Type 2 Diabetes. J Gerontol A Biol Sci Med Sci 71, 850–857 (2016).

60. Suriano, F. et al. Novel insights into the genetically obese (ob/ob) and diabetic (db/db) mice: two sides of the same coin. Microbiome 9, 147 (2021).

61. Yahagi, N. et al. p53 Involvement in the Pathogenesis of Fatty Liver Disease*. Journal of Biological Chemistry 279, 20571–20575 (2004).

62. Kracikova, M., Akiri, G., George, A., Sachidanandam, R. & Aaronson, S. A. A threshold mechanism mediates p53 cell fate decision between growth arrest and apoptosis. Cell Death Differ 20, 576–88 (2013).

63. Soto-Gutierrez, A., Gough, A., Vernetti, L. A., Taylor, D. & Monga, S. P. Pre-clinical and clinical investigations of metabolic zonation in liver diseases: The potential of microphysiology systems. Exp Biol Med 242, 1605–1616 (2017).

64. Chandel, N. S., Vander Heiden, M. G., Thompson, C. B. & Schumacker, P. T. Redox regulation of p53 during hypoxia. Oncogene 19, 3840–3848 (2000).

65. Koumenis, C. et al. Regulation of p53 by Hypoxia: Dissociation of Transcriptional Repression and Apoptosis from p53-Dependent Transactivation. Mol Cell Biol 21, 1297–1310 (2001).

66. Postic, C. et al. Dual Roles for Glucokinase in Glucose Homeostasis as Determined by Liver and Pancreatic β Cell-specific Gene Knock-outs Using Cre Recombinase. Journal of Biological Chemistry 274, 305–315 (1999).

67. Marino, S., Vooijs, M., van Der Gulden, H., Jonkers, J. & Berns, A. Induction of medulloblastomas in p53-null mutant mice by somatic inactivation of Rb in the external granular layer cells of the cerebellum. Genes Dev 14, 994–1004 (2000).

68. Hock, A. K. et al. Development of an inducible mouse model of iRFP713 to track recombinase activity and tumour development in vivo. Sci Rep 7, 1837 (2017).

69. Bankhead, P. et al. QuPath: Open source software for digital pathology image analysis. Sci Rep 7, 16878 (2017).

70. Humpton, T. J., Hock, A. K., Maddocks, O. D. K. & Vousden, K. H. p53-mediated adaptation to serine starvation is retained by a common tumour-derived mutant. Cancer Metab 6, 18 (2018).

71. Luo, Y., Rana, P. & Will, Y. Palmitate increases the susceptibility of cells to drug-induced toxicity: An In Vitro method to identify drugs with potential contraindications in patients with metabolic disease. Toxicological Sciences 129, 346–362 (2012).

72. Humpton, T. J. et al. Oncogenic KRAS Induces NIX-Mediated Mitophagy to Promote Pancreatic Cancer. Cancer Discov 9, 1268–1287 (2019).

