## Supplemental Materials for "p53 and TIGAR promote redox control to protect against metabolic dysfunction-associated steatohepatitis"

*Supplemental Figure 1, Wittke et al.*

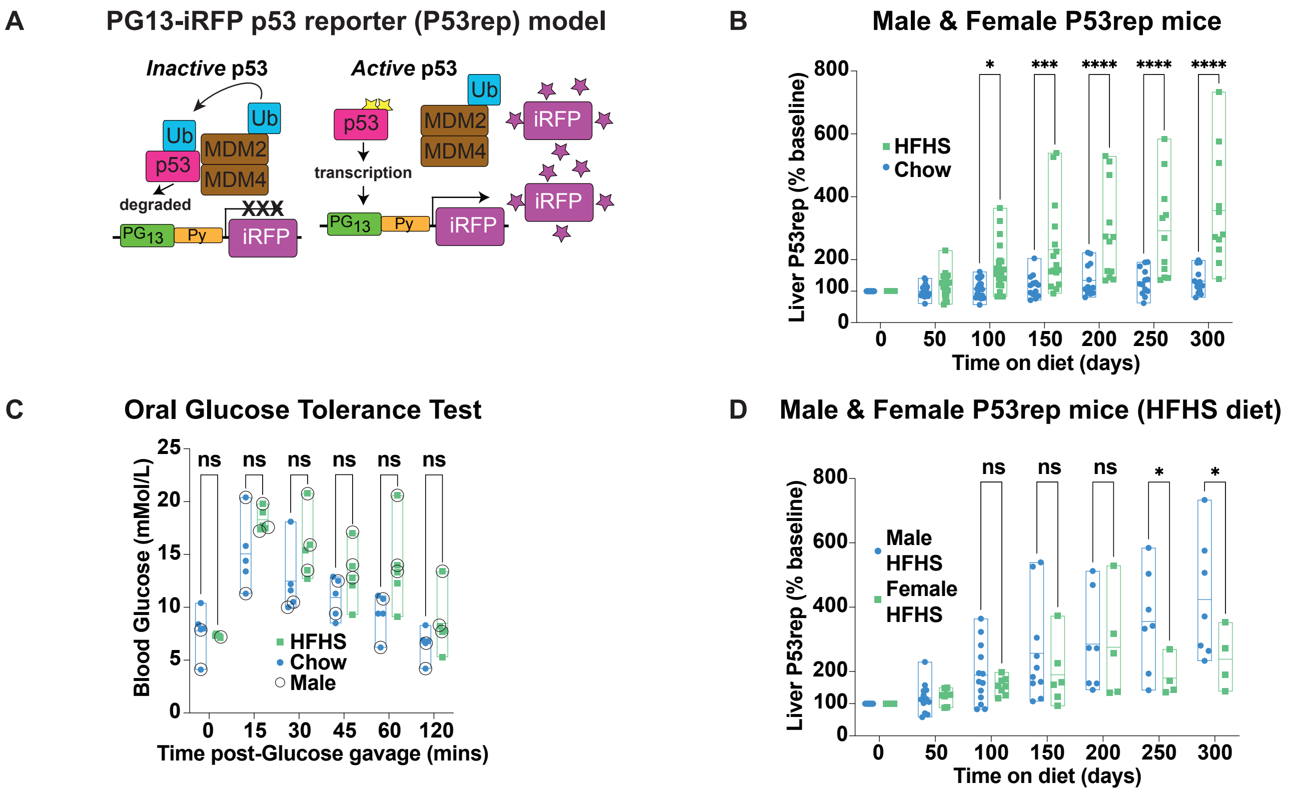


**Supplemental Figure 1: Liver p53 is engaged more strongly in male mice in response to a high fat and high sugar diet**

**A.** Model of how activation of the p53 pathway is monitored using the PG13-iRFP p53 (P53rep) reporter. P53 is constitutively expressed but rapidly degraded. Inactive p53 is turned over via MDM2-MDM4-mediated ubiquitination (Ub) and subsequent proteasomal degradation. Activation of p53 occurs via post-translational modifications which promote the stability of p53. This leads to the transcription of p53 target genes (those containing p53-binding sequences, including the PG sequence used in the P53rep construct). In P53rep mice, iRFP expression occurs in response to active p53 signalling and can be non-invasively monitored over time.

**B.** Pooled quantification of liver-region P53rep signal in male and female mice as shown in Figure 1 (A-D). Quantification is normalised to the initial liver signal identified per mouse. N=21 chow and N=21 HFHS mice at 0d, 50d, and 100d diet; N=13 chow and N=16 HFHS mice at 150d diet; N=13 chow and N=12 HFHS mice at 200d diet; and N=13 chow and N=11 HFHS mice at 250d and 300d diet. Data presented as mean +/- range with individual data points shown. Data analysed using a Two-way ANOVA with Holm-Sidak's multiple comparisons test and multiplicity-adjusted p-values. *p<0.05, ***p<0.001, ****p<0.0001.

**C.** Oral glucose tolerance test performed in P53rep mice after 100 days on either HFHS or control (chow) diet. Blood glucose levels (in mMol/L) were measured at the indicated intervals (minutes) after administration of glucose gavage. N=5 mice per condition with N=2 male and N=3 female mice in the chow group and N=3 male and N=2 female mice in the HFHS group. Data presented and analysed as in (B). ns- not significant. Male mice in each cohort marked with circles.

**D.** Quantification of liver-region P53rep signal in male and female mice on the HFHS diet as shown in Figure 1 (A-D). Quantification is normalised to the initial liver signal identified per mouse. N=13 male and N=8 female HFHS mice at 0d, 50d, and 100d diet; N=10 male and N=6 female HFHS mice at 150d diet; N=7 male and N=5 female HFHS mice at 200d diet; and N=7 male and N=4 female HFHS mice at 250d and 300d diet. Data presented and analysed as in (B). ns- not significant, *p<0.05.

### *Supplemental Figure 2, Wittke et al.*

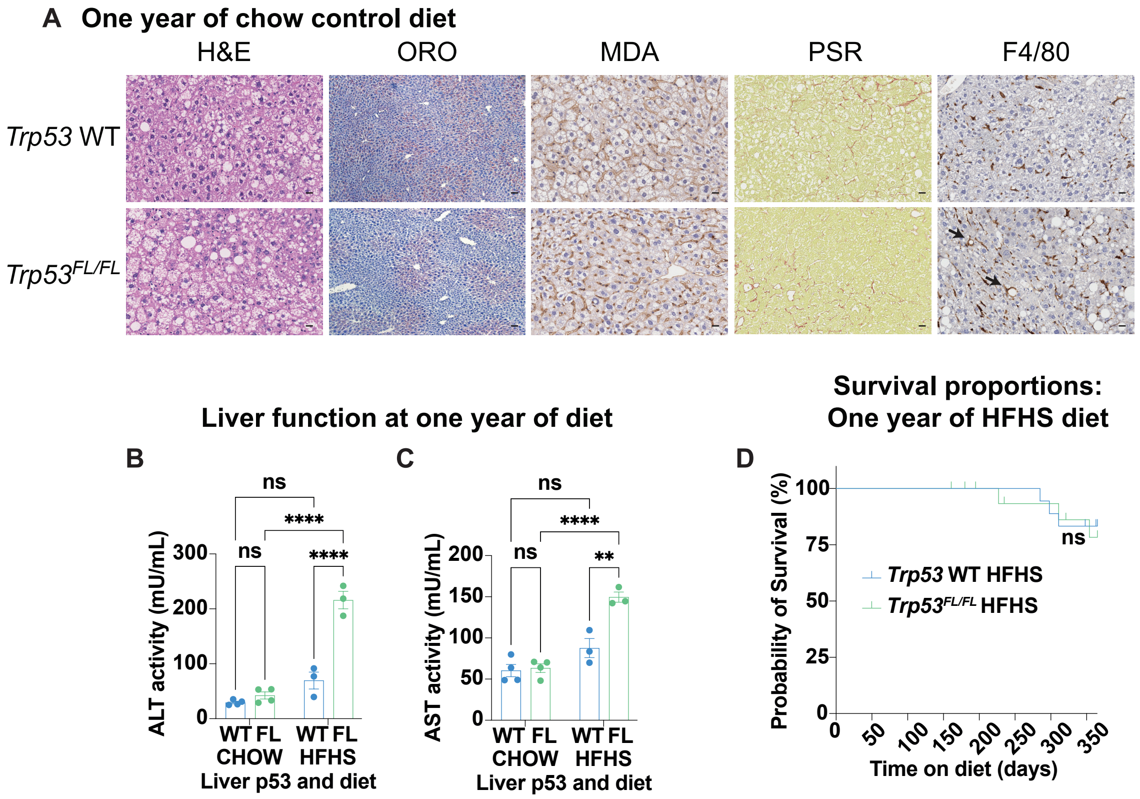


**Supplemental Figure 2: Liver p53 limits liver damage arising from a high fat and high sugar diet *in vivo***

**A.** Representative H&E images, staining for lipid content (oil-red-o / ORO), IHC staining lipid peroxidation (malondialdehyde/MDA), staining for fibrosis (picrosirius red/PSR), and IHC staining for macrophages (F4/80) in male *Albumin*-Cre+; *Trp53* WT and *Albumin*-Cre+; *Trp53^FL/FL^* (liver-specific deletion of *Trp53*) (*Trp53^FL/FL^*) mice after one year on an obesogenic high fat and high sugar (HFHS) diet. For ORO images, representative of N=4 *Trp53* WT chow-fed mice, N=3 *Trp53^FL/FL^* chow-fed mice and N=3 HFHS-fed mice per genotype; scale bars 40 μm. For the remainder, representative of N=9 *Trp53* WT chow-fed mice, N=8 *Trp53^FL/FL^* chow-fed mice and N=13 HFHS-fed mice per genotype. Scale bars 20 μm. Arrows denote F4/80-positive lipogranulomas indicative of chronic inflammation.

**B/C.** Plasma activity (mU/mL) of alanine (B) and aspartate (C) transaminase (ALT/AST) to assess liver function in male *Trp53* WT and *Trp53^FL/FL^* (FL) mice after one year on an obesogenic high fat and high sugar (HFHS) diet compared with aged-matched mice on normal diet (chow). N=4 mice/chow-fed group and N=3 mice/HFHS-fed group. Each data point represents the mean from technical duplicates per mouse. Data presented as mean +/- SEM and analysed using two-way ANOVA with Holm-Sidak's multiple comparisons test and multiplicity-adjusted p-values. ns- not significant. **p<0.01, ****p<0.001.

**D.** Survival percentages for one year on the HFHS diet within male *Albumin*-Cre+; *Trp53* WT and *Albumin*-Cre+; *Trp53^FL/FL^* mice shifted onto a HFHS diet at 65-75 days of age. N=18 mice / group. Data analysed using the Log-rank (Mantel-Cox) test. ns- not significant.

*Supplemental Figure 3, Wittke et al.*

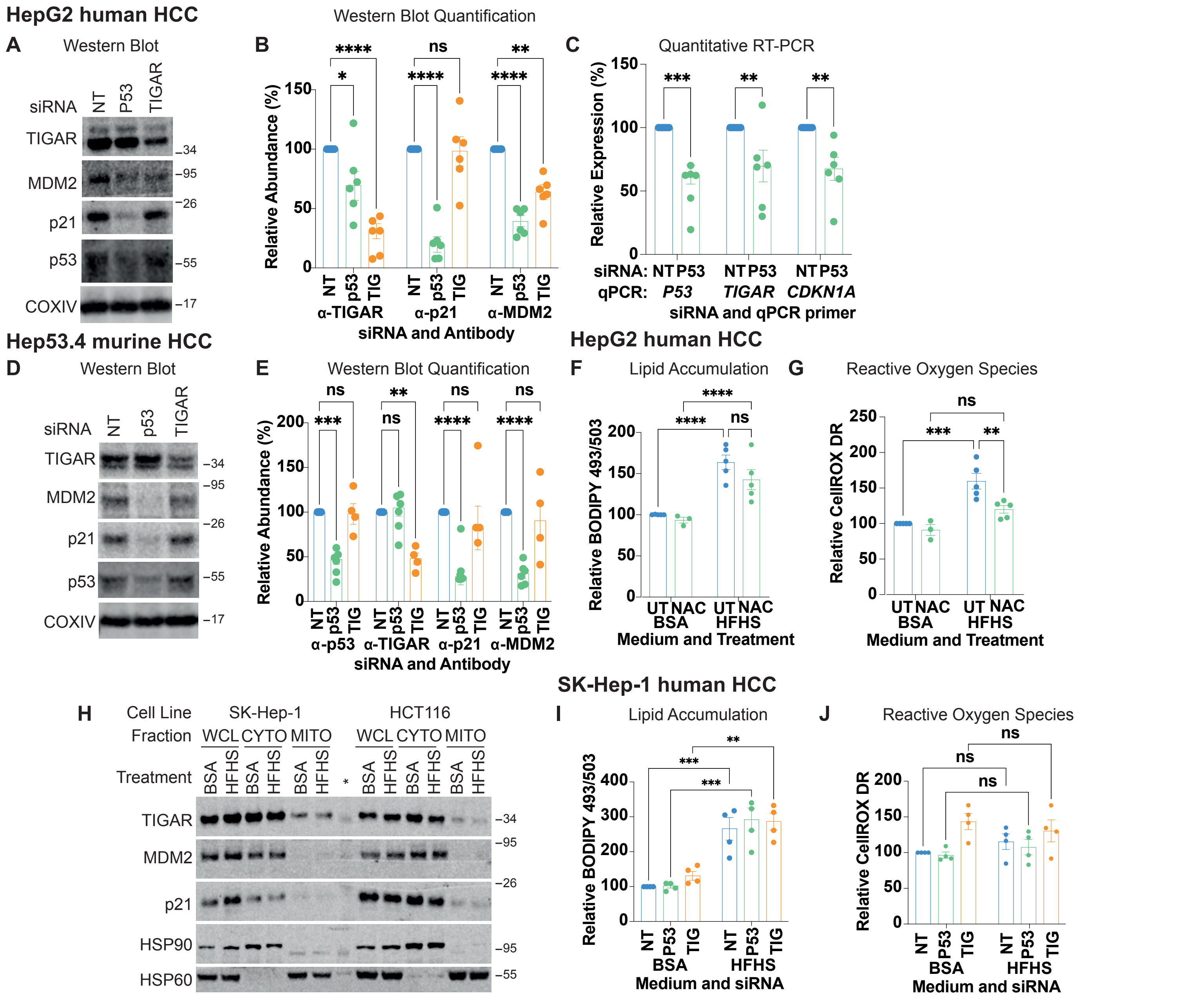


**Supplemental Figure 3: p53 supports redox control and engages TIGAR in p53 WT HepG2 and Hep53.4 HCC cell lines in response to overfeeding *in vitro***

**A/B.** Western blot analysis (A) and quantification (B) of TIGAR, MDM2, and CDKN1A/p21 in p53 WT HepG2 cells treated with either non-targeting control (NT) or siRNA directed against TP53 (P53) or TIGAR for 96 hours. Representative of N=5 independent experiments. COXIV used as loading control. TIGAR abundance represents the sum of bands. Protein MW ladder (in kilodaltons) as depicted. Data presented as mean +/- SEM with data points and analysed by two-way ANOVA with Holm-Sidak's multiple comparisons test and multiplicity-adjusted p-values: ns- not significant, *p<0.05, **p<0.01, ****p<0.0001.

**C.** Quantitative RT-PCR analysis of P53, TIGAR, and CDKN1A/p21 expression in p53 WT HepG2 cells treated with either non-targeting control (NT) or siRNA directed against TP53 (P53) for 96 hours. Representative of N=6 independent experiments. Data presented and analysed as (B): **p<0.01, ***p<0.001.

**D/E.** Western blot analysis (D) and quantification (E) of TIGAR, MDM2, and CDKN1A/p21 expression in Trp53 WT Hep53.4 cells treated with either non-targeting control (NT) or siRNA directed against Trp53 (P53) or Tigar (TIG) for 72 hours. Representative of N=6 independent experiments for NT/Trp53 siRNA and N=4 for Tigar siRNA. COXIV used as loading control. Protein MW ladder (in kilodaltons) as depicted. Data presented and analysed as in (B). ns- not significant, **p<0.01, ***p<0.001, ****p<0.0001.

**F/G.** Analysis of lipid content (BODIPY 493/503) (F) or ROS (CellROX Deep Red) (G) via flow cytometry in *TP53* KO HepG2 cells after 48 hours of culture in either control (BSA) or HFHS medium supplemented with PBS control (UT) or 2 mM N-acetyl-cysteine (NAC). Data from N=5 experiments with HFHS+NAC, of which only N=3 experiments also included BSA+NAC. Data presented and analysed as in (B): ns- not significant, **p<0.01, ***p<0.001, ****p<0.0001.

**H.** Western blot analysis of TIGAR, MDM2, and CDKN1A/p21 expression in p53 WT SK-Hep-1 and HCT116 cells grown in baseline medium supplemented with BSA (BSA) or with a high fat and high sugar (HFHS) medium formulation for 48 hours. Whole cell lysate (WCL), Cytoplasmic (CYTO) and mitochondrial-enriched fractions (MITO) as indicated. *- protein ladder. HSP90 used as cytoplasmic loading control and COXIV used as mitochondrial loading control. Protein MW ladder (in kilodaltons) as depicted. Representative of N=3 independent experiments per cell line.

**I/J.** Analysis of lipid content (BODIPY 493/503) (I) and ROS (CellROX Deep Red) (J) via flow cytometry in p53 WT SK-Hep-1 cells treated with either non-targeting control (NT) or siRNA directed against TP53 (P53) or TIGAR (TIG) for 96 hours and cultured in either baseline medium (supplemented with BSA) or HFHS medium for the final 48 hours prior to analysis. Data from N=4 experiments and presented as median fluorescence intensity (MFI) relative to NT baseline condition per experiment. Data presented and analysed as in (B), ns- not significant, **p<0.01, ***p<0.001.

*Supplemental Figure 4, Wittke et al.*

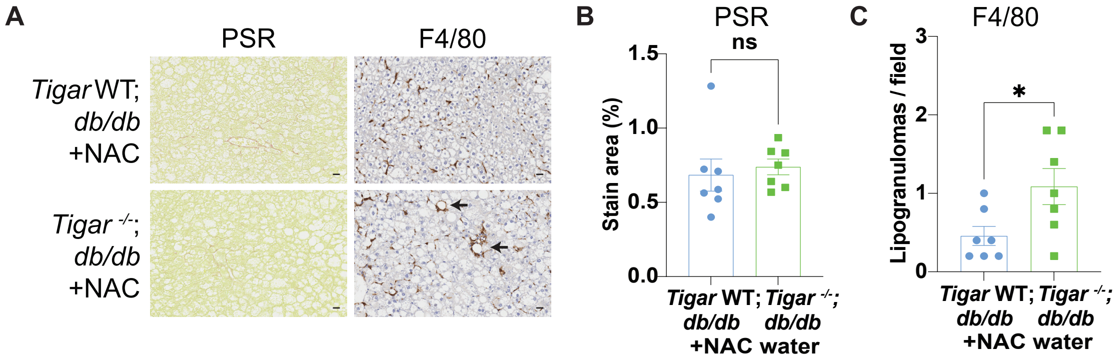


**Supplemental Figure 4: N-acetyl-cysteine supplementation reduces ROS in leptin receptor-deficient (*Lepr*$^{db/db}$) mice that lack *Tigar* expression**

**A.** Representative staining for fibrosis (picrosirius red/PSR) and IHC staining for macrophages (F4/80) in female *Lepr*$^{db/db}$ *(db/db)* (*Tigar* WT) and *Tigar*$^{-/-}$; *db/db* mice at 100 days of age and treated with N-Acetyl-Cysteine (NAC) in the drinking water from 42 days of age. Representative of N=7 *db/db* (*Tigar* WT) and N=7 *Tigar*$^{-/-}$; *db/db* mice. Macrophage-engulfed steatotic hepatocytes (lipogranulomas) depicted by arrows in F4/80 images. Scale bars 20 µm.

**B/C.** Quantification of PSR to assess fibrosis (B) and F4/80 to assess lipogranulomas (C) in *db/db* (*Tigar* WT) and *Tigar*$^{-/-}$; *db/db* mice at 100 days of age, treated with N-Acetyl-Cysteine (NAC) in the drinking water from 42 days of age (as in A). N=7 *db/db* (*Tigar* WT) + NAC and N=7 *Tigar*$^{-/-}$; *db/db* mice + NAC. Data presented as mean +/- SEM and analysed using an unpaired t-test with Welch's correction. ns- not significant, *p<0.05.
